# Separable cellular correlates of visual acuity and lifespan in mammalian primary visual cortex

**DOI:** 10.64898/2026.04.08.716558

**Authors:** Daniel J. Miller, Jon H. Kaas

## Abstract

Visual acuity spans more than a 100-fold range in mammals and yet the neural correlates of this perceptual gradient has not been fully evaluated. Furthermore, even though the known derived features of the human brain include specific changes to the cerebral cortex and the visual system in particular, no evolution of developing life histories approach has been quantitatively applied to metrics of cell composition. In this study, we present stereological estimates of neuron and glia density in V1 and granular layer 4 in a comparative sample of primates. We then integrate these data with the literature to construct a larger comparative dataset to test for phylogenetic relationships in mammalian visual system organization. Our examination revealed a primary relationship between acuity and neuron number along with secondary relationships among cell types tied to metabolic maintenance and support across the lifespan. Retinal metabolics along with phylogenetic position accounts for a large amount of V1 neuron density, which are further related to acuity while glia are related to longevity. These results identify a dissociation in the evolutionary developmental organization and senescence of V1 that map onto first principal parameters to explore the phylogenetic position and ecological pressures acting on the mammalian visual system. In accord with the literature, humans are revealed as outliers in glial support of neuronal metabolism across the lifespan. These findings provide evidence that mammalian visual cortex varies along at least two partially separable cellular dimensions in which visual resolution differs from lifelong maintenance.

**Summary and Significance:** If the cellular organization of visual cortex is shaped by a single constraint or multiple independent pressures is debated. We used stereology to test whether V1 neurons and glia make separable contributions to visual performance across the mammal lifespan. V1 neuron density, together with brain size, predicts visual acuity. The glia-to-neuron ratio is alternatively associated with maximum lifespan after controlling for neuron density. Humans and chimpanzees have nearly identical V1 neuron densities, yet humans appear show substantially elevated glial investment across the literature. These findings suggest that mammalian visual cortex evolves under at least two partially separable pressures with neuron density and spatial resolution on one hand and glia investment to sustain function across life on the other.

## Introduction

Primary visual cortex (V1) has occupied a central place in neuroscience since the Stria of Gennari in layer 4 was suggested as an early defining anatomical landmark by which this cortical field could be identified across species (1–4). Its functional importance was established later by the physiological work of Hubel and Wiesel (5,6), who showed that V1 neurons transform retinal input into organized signals for orientation, spatial frequency, and ocular dominance (7,8). Subsequent work connected this physiological organization to visual performance through the cortical magnification factor (9), demonstrating that the amount of cortex devoted to representing the retinal image predicts psychophysical resolution (10). Together, these traditions established a broad principle that visual capability depends not only on the properties of the eye, but also on the amount and organization of cortical tissue allocated to processing visual input (11–13). From that foundation, alternative literatures developed with the first studies of how cortical cell organization scales with peripheral input, including how neuronal number and packing vary with retinal and thalamic afferent structure across species (8,12–20). Others examined the energetic cost of operating visual circuits and the selective pressures that metabolic expense imposes on visual-system design, including lifelong function (11,21–26). Work often proceeds in parallel and thus we still are not sure if interspecific variation in V1 cellular composition predicts visual acuity directly or is related to some other first principle of homeostatic restoration (27).

Comparative work on V1 cellular organization has focused primarily on how neuron number scales across species and within the brain (12,14,16,19). Kaskan et al. (15) argued that central visual structures scale relatively conservatively with brain size, implying that major visual specialization is expressed chiefly at the retinal level. By contrast, Herculano-Houzel and colleagues (19,28) showed that neuron density varies substantially across cortical areas and among primate species to suggest that cortical organization is relatively modifiable. Stevens (12) then provided a formal account of one such pattern by showing that V1 neuron number scales as the 3/2 power of LGN neuron number across haplorhine primates, a geometric consequence of mapping a two-dimensional retinal surface into a three-dimensional cortical representation. Srinivasan et al. (13) extended this structural result into a functional prediction via the Nyquist-sampling logic where denser neuronal packing should support finer spatial resolution of the visual field. What remains untested is whether that prediction holds across a broader mammalian sample and if cell types differentially contribute to acuity.

An energetic tradition approached visual-system design through the metabolic cost of neural signaling (21). The Attwell and Laughlin tradition showed that Na⁺/K⁺-ATPase-mediated restoration of ionic gradients after synaptic transmission accounts for a major share of cortical energy expenditure across the visual pathway (23,29), and suggest the broader evolutionary implication that energy limitation acts as a pervasive selective pressure on sensory-system design (11). Most relevant here, Heras and Laughlin recently (11) formalized a photoreceptor energy tariff and showed that the optical parameters governing spatial resolution are relatively insensitive to that tariff, whereas the metabolic depth of the photoreceptor array that can be sustained is highly sensitive to it. This dissociation between determinants of performance and determinants of maintenance affordability (25,26) motivates a parallel prediction at the cortical level (28,30) where V1 neuron density should predict acuity and the GNR, as an index of support burden rather than sampling resolution, should not.

Glia provide the cellular infrastructure through which cortex manages the metabolic and homeostatic costs of neural signaling (31), making GNR a plausible index of maintenance investment per unit of computational capacity (28,30). Astrocytes regulate neurotransmitter recycling, extracellular ions, and neurovascular coupling, whereas oligodendrocytes reduce conduction cost through activity-dependent myelination while also imposing a continuing maintenance burden (32,33). Across species, however, GNR may also scale inversely with neuronal density, with smaller and more densely packed neurons requiring less glial support per cell (34–36). Residual variation above or below that structural baseline is therefore potentially informative. In that context, the elevated GNR reported in humans relative to other hominoids in frontal cortex (30) and relative to the nonhuman primate V1 distribution these data suggest that human glial enrichment (37–39) may reflect a broader evolutionary pattern.

The prediction that residual GNR should covary with lifespan rather than acuity draws on classical actuarial and life-history theory (26). Just over 200 years ago, Gompertz showed that mortality increases exponentially with age, establishing aging as a process with independently variable temporal parameters (25). Sacher later argued that interspecific differences in longevity arise primarily from differences in the slope of the log-mortality curve, that is, in the rate at which biological maintenance deteriorates (24). Hou (40) provided an energetic extension of this framework by decomposing the Gompertz growth constant into mass-specific metabolic rate and biosynthetic cost, and showed that biosynthetic cost predicts maximum lifespan independently of metabolic rate across mammals. These results imply that investment in construction and maintenance of the phenotype is selectable independently of operational throughput (41,42). If glia instantiate that maintenance investment in cortex (30), then GNR, after accounting for neuronal packing geometry, should covary with maximum lifespan, whereas neuron density should index a separable performance axis (11,13,15,16).

Here we test the idea of developmental biosynthetic parameterization in a 15 species, comparative dataset that combines original stereological measurements from nine primate genera with published data on retinal energetics, thalamocortical architecture, V1 cellular composition, visual acuity, and maximum lifespan to detect phylogenetic relationships (11–16,19–21,23,37–39). To specifically clarify the relationship among ecological light levels, visual performance, and neural organization, we studied the cellular composition of the visual system. To our knowledge, this is the largest sample of V1 thalamorecipient layer 4 cell counts investigated in primates to date.

## Results

We obtained design-based stereological estimates of V1 volume and cell composition for nine primate genera spanning the basal prosimian galago (*Otolemur garnettii*) to humans (*Homo sapiens*) using Cavalieri’s method for volumetrics and the optical fractionator (35,43) for cell counts (Figure 1, Table 1; Figures S1–S8).

**Figure 1.**
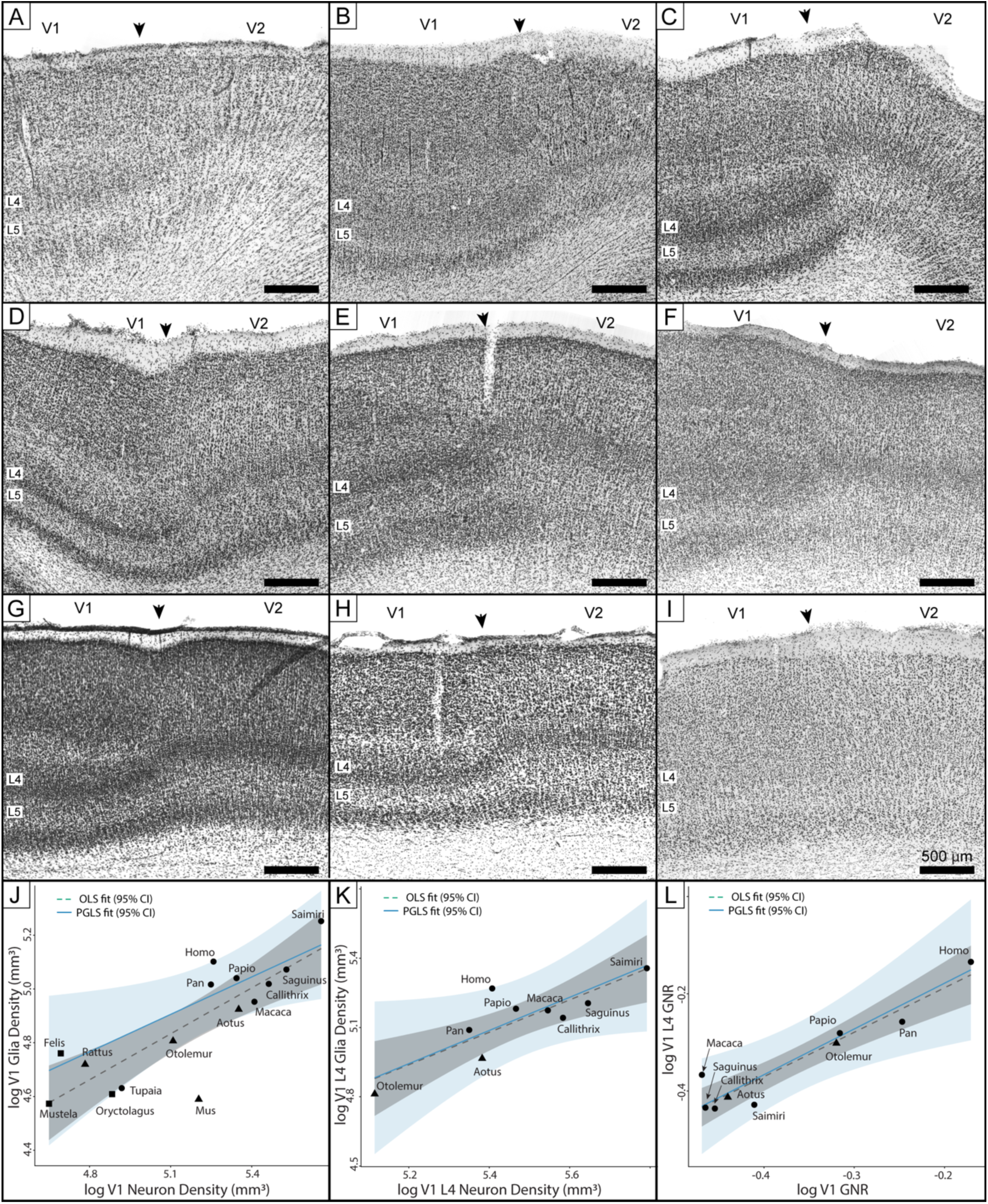
Cellular organization of primate V1. Panels A to I show low magnification images of Nissl-stained sections at the boundary between the primary (V1) and secondary visual cortical fields (V2) for the nine primate species studied. Arrowheads mark the cytoarchitectonic boundary between V1 and V2, identifiable by modulations in the cellular composition of layer 4 (L4) across the transition (4), with layer 5 (L5) delineated for comparison. Scale bar represents 500 μm and applies to all panels A–I. Panels J–L show relationships between neuron and glial cell densities in V1 and layer 4, estimated by design-based stereology for the nine primate species shown in panels A–I. In all three panels, solid lines depict OLS regression fits with 95% confidence intervals (shaded), and dashed lines depict PGLS regression fits with approximate confidence intervals (shaded). Each species is labeled by genus name.

**Table 1.**
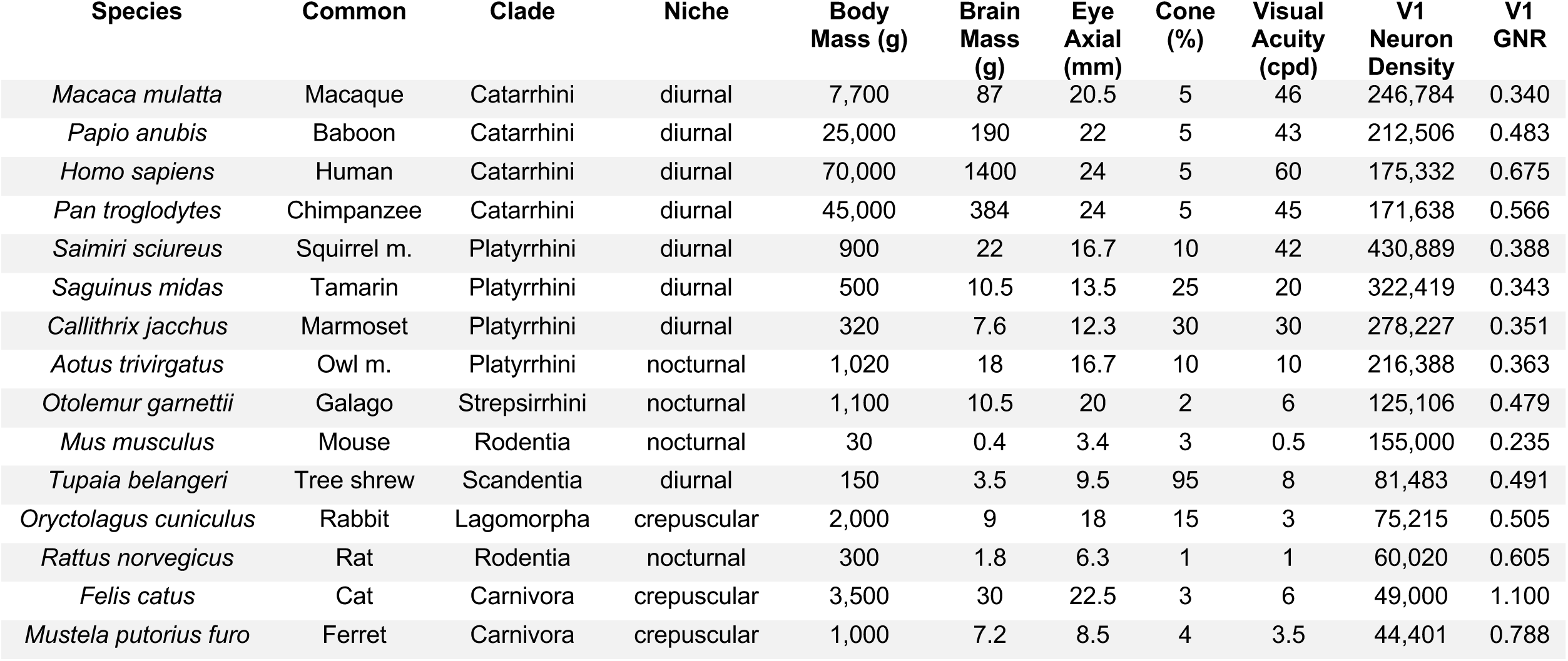
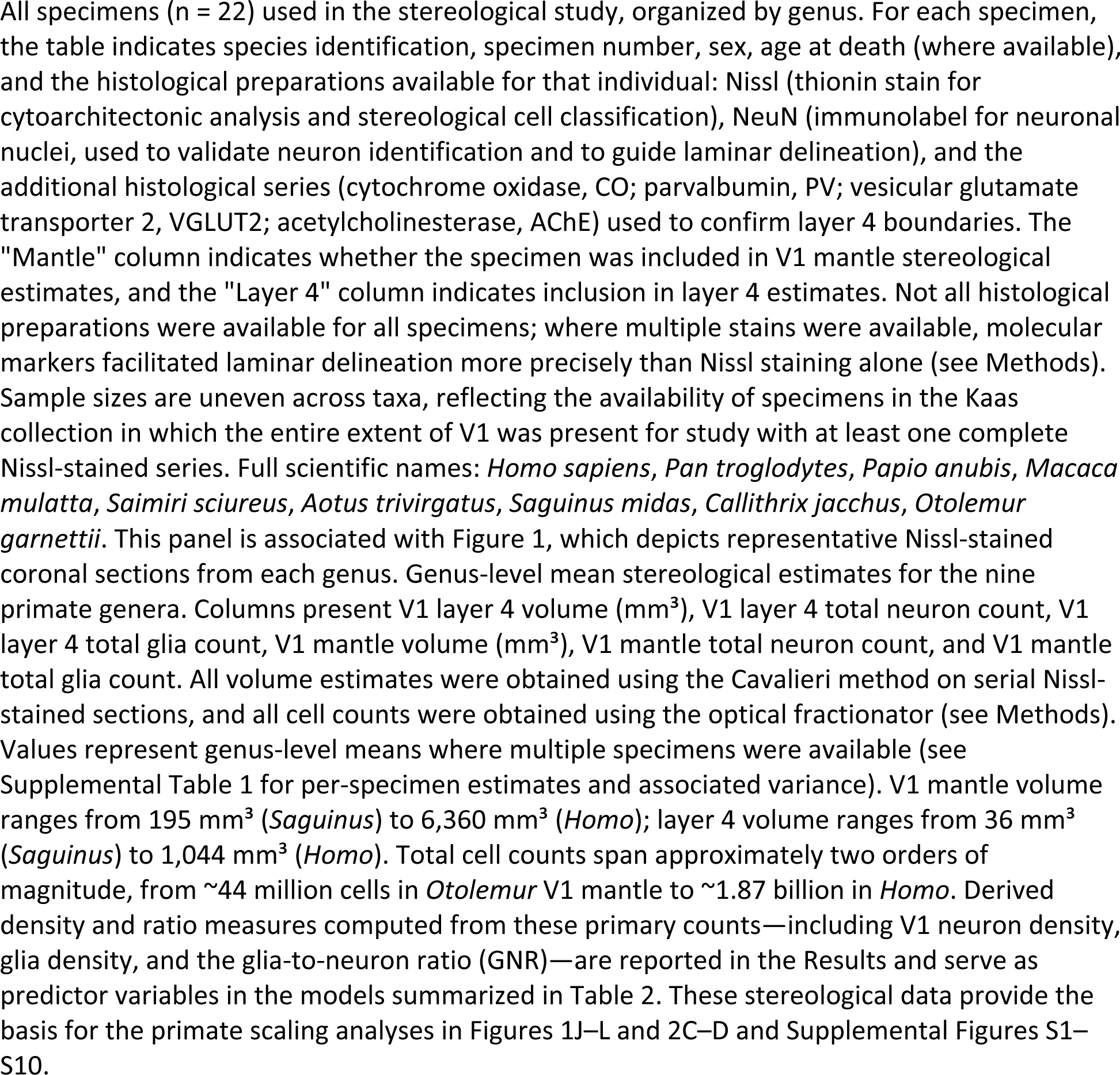
Specimens and data.

Volumetrics vary more than 30-fold across the sample. Whole V1 volumes range from 195 mm³ in the tamarin (*Saguinus*) to 6,360 mm³ in humans (*Homo*), and layer 4 volumes range from 36 mm³ to 1,044 mm³, respectively (Table 1). Despite this absolute expansion, the volumetric proportion of layer 4 within V1 is broadly conserved, averaging 19.3% (SD = 2.2%). *Aotus* (22.8%) and *Macaca* (22.6%) show the highest layer 4 proportions while humans shows the lowest (16.4%). Total V1 cell number spans more than 40-fold and cell number varies from approximately 44.4 million cells in galagos (*Otolemur*) to approximately 1.87 billion cells in humans (*Homo*). Neuron density in the V1 mantle and layer 4 are highest in diurnal platyrhines followed by catarhines at an intermediate position followed by nocturnal strepsirhines (Table 1).

Glia scale positively with neurons per unit volume. Specifically, the density of cell types scale in both whole V1 and layer 4 (Figure 1, Table 2, Figures S9–S12, Table S1), exhibiting near-isometric PGLS slopes (0.94–0.95) and strong phylogenetic signal (Pagel’s λ ≈ 1.0). The GNR in the V1 mantle ranges from 0.340 in *Macaca* to 0.675 in *Homo*, with a nine-genus mean of 0.443 and a nonhuman primate mean of 0.414. Human GNR exceeds the sampled primate mean by more than 50% (Figure 1, Table 1, Table 2), both across V1 and within layer 4 (Figures S9-S12, Table S1).

**Table 2.**
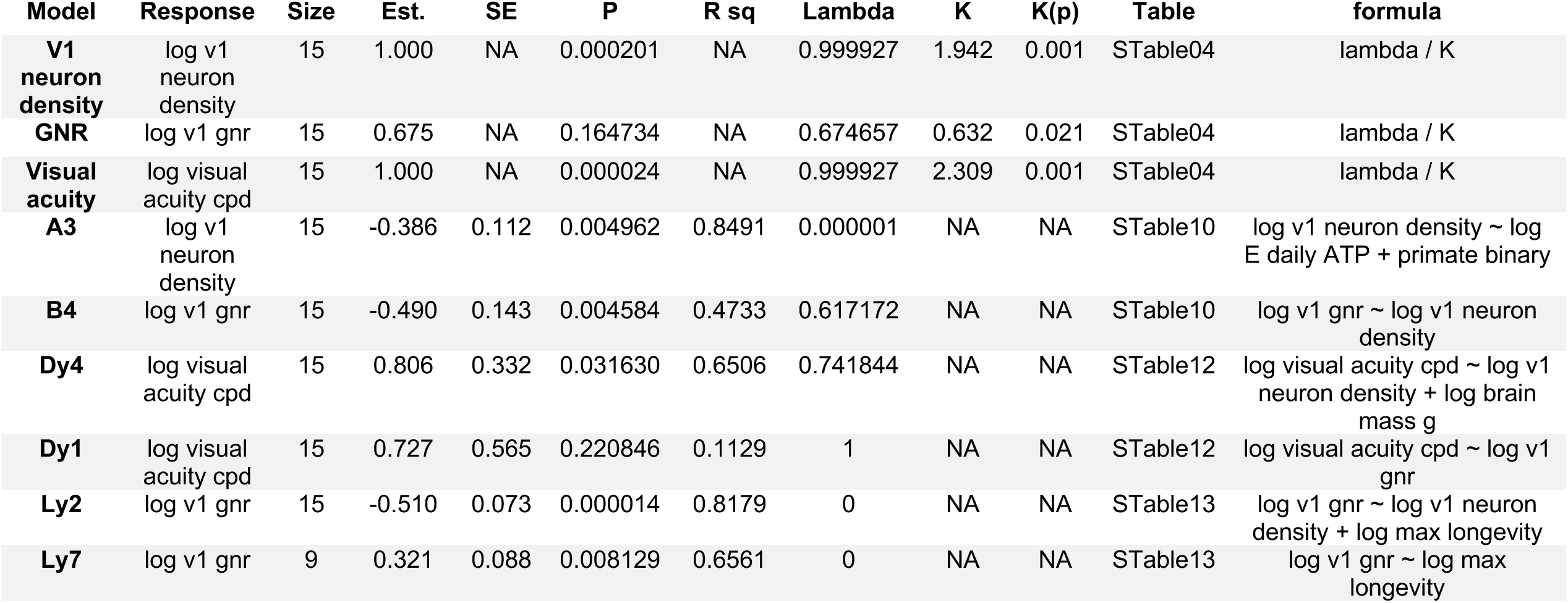
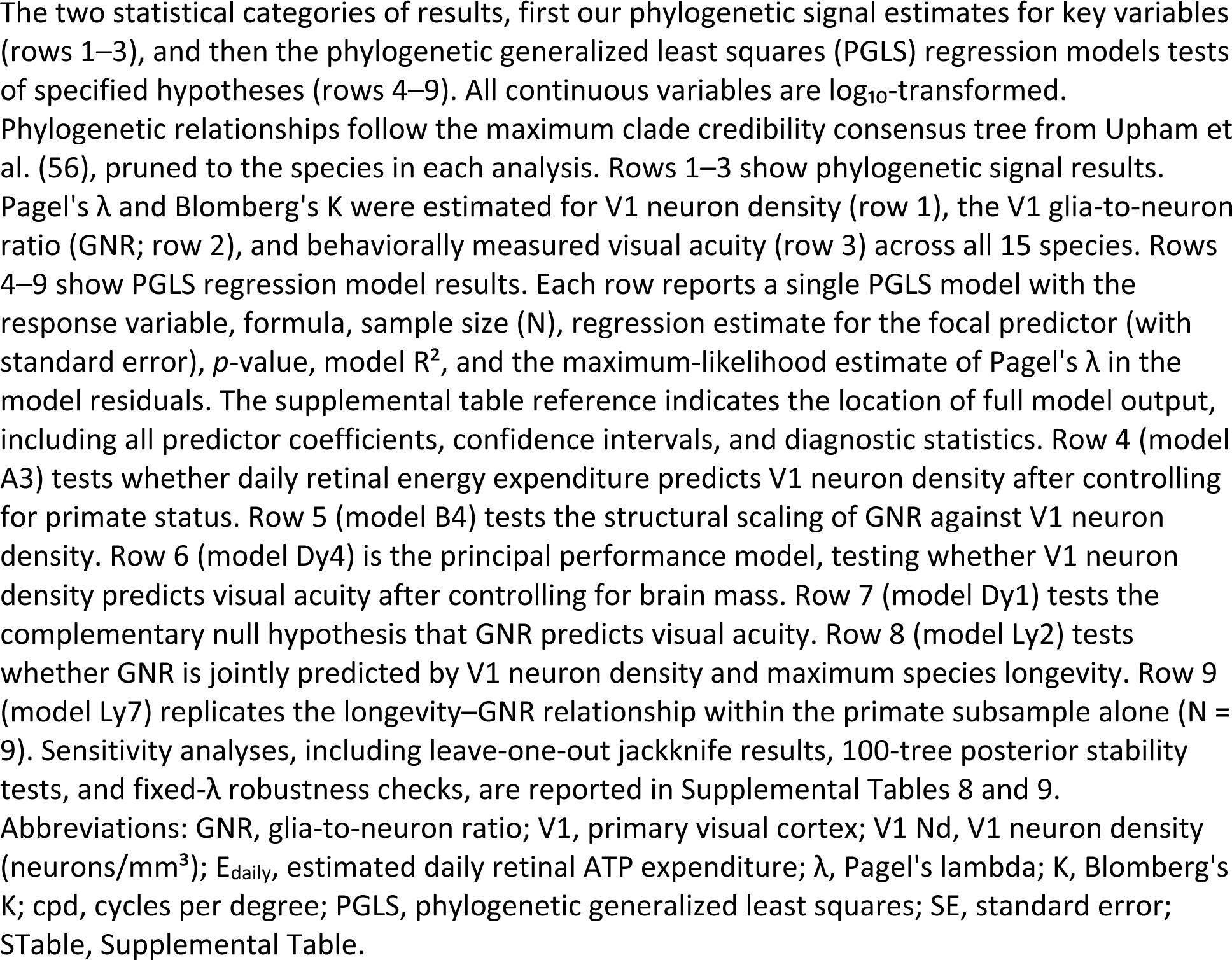
Summary of phylogenetic signal and principal model result.

We surveyed the literature on the visual system (16) to make a larger mammalian sample at 15 taxa to test hypotheses (Figure 2, Table 1). Visual pathway cell counts (retina to LGN to V1) and a composite magnification index summarizing total V1 cellular investment per retinal ganglion cell fiber are reported for the nine primate genera in (Figure 2, Figure 3, Table S2). V1 neuron density and visual acuity each carry strong phylogenetic signal across the 15-species dataset with Pagel’s λ = 1.000 and Blomberg’s K > 1.9 for both variables (p ≤ 0.001; Table 2). GNR shows weaker signal at λ = 0.675 (p = 0.165, non-significant), though Blomberg’s K is significant (K = 0.632, p = 0.037, Table 2). The composite transform index scales positively with brain mass across primates (Figure 2, Table S2), consistent with larger-brained species routing more total cellular investment through the primary visual pathway per peripheral fiber.

**Figure 2.**
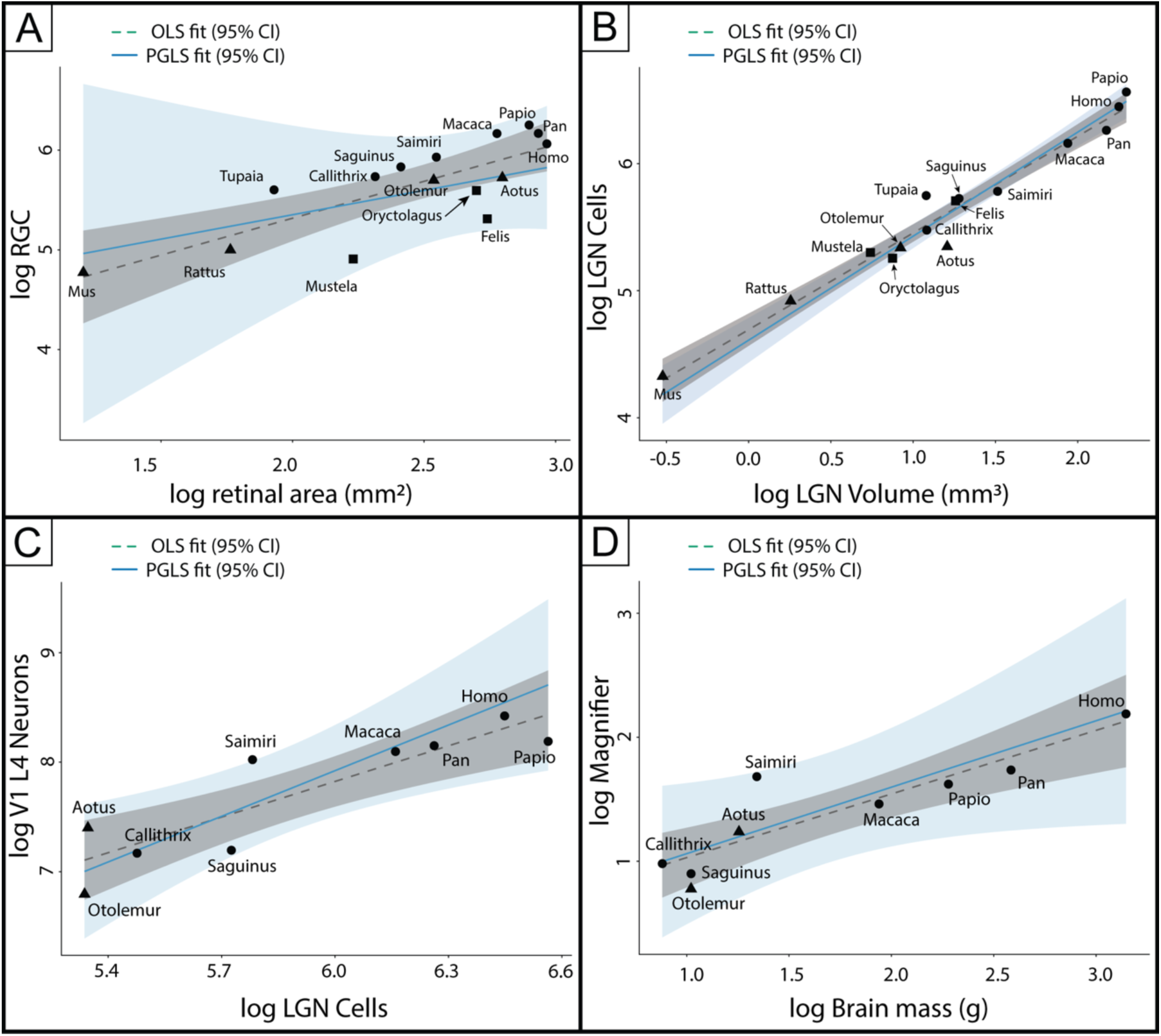
Cellular scaling from the primate retina to cortex. Four panels trace the quantitative relationships among cell populations at successive stages of the primary visual pathway from the retina to LGN to V1 layer 4 and overall brain mass. Panel (A) shows retinal ganglion cell (RGC) count as a function of retinal area (mm²) across 15 mammalian species spanning five orders. Panel (B) shows LGN neuron count as a function of LGN volume (mm³) across all 15 species. Panel (C) shows V1 layer 4 total neuron count as a function of LGN neuron count for the nine primate genera, representing the thalamocortical expansion from relay nucleus to principal cortical input layer. Panel (D) shows the log magnifier term as a function of log brain mass (g) for the nine primate genera. The magnifier term is defined as T1 × T2 × (1 + GNR), where T1 is the retinogeniculate transform (LGN neurons per RGC), T2 is the geniculocortical transform (V1 layer 4 neurons per LGN neuron), and GNR is the indexed V1 GNR. This composite metric estimates the total effective cellular investment in V1 layer 4 per retinal ganglion cell fiber, incorporating both the computational substrate and the metabolic overhead of the associated glial population (see Figure 3 for the full circuit-level decomposition). All axes are log₁₀-transformed, and in each panel, dashed green lines represent ordinary least-squares (OLS) regression fits and solid blue lines represent phylogenetic generalized least-squares (PGLS) fits, each with 95% confidence intervals (grey and blue shading, respectively). Species are labeled by genus name, and symbol shapes denote taxonomic grouping (filled circles, primates; filled triangles, non-primate mammals with literature-derived counts; filled squares, species for which values were estimated, see Methods). All 15 species are shown in panels A and B, while panels C and D are restricted to the nine primate genera with original stereological data.

**Figure 3.**
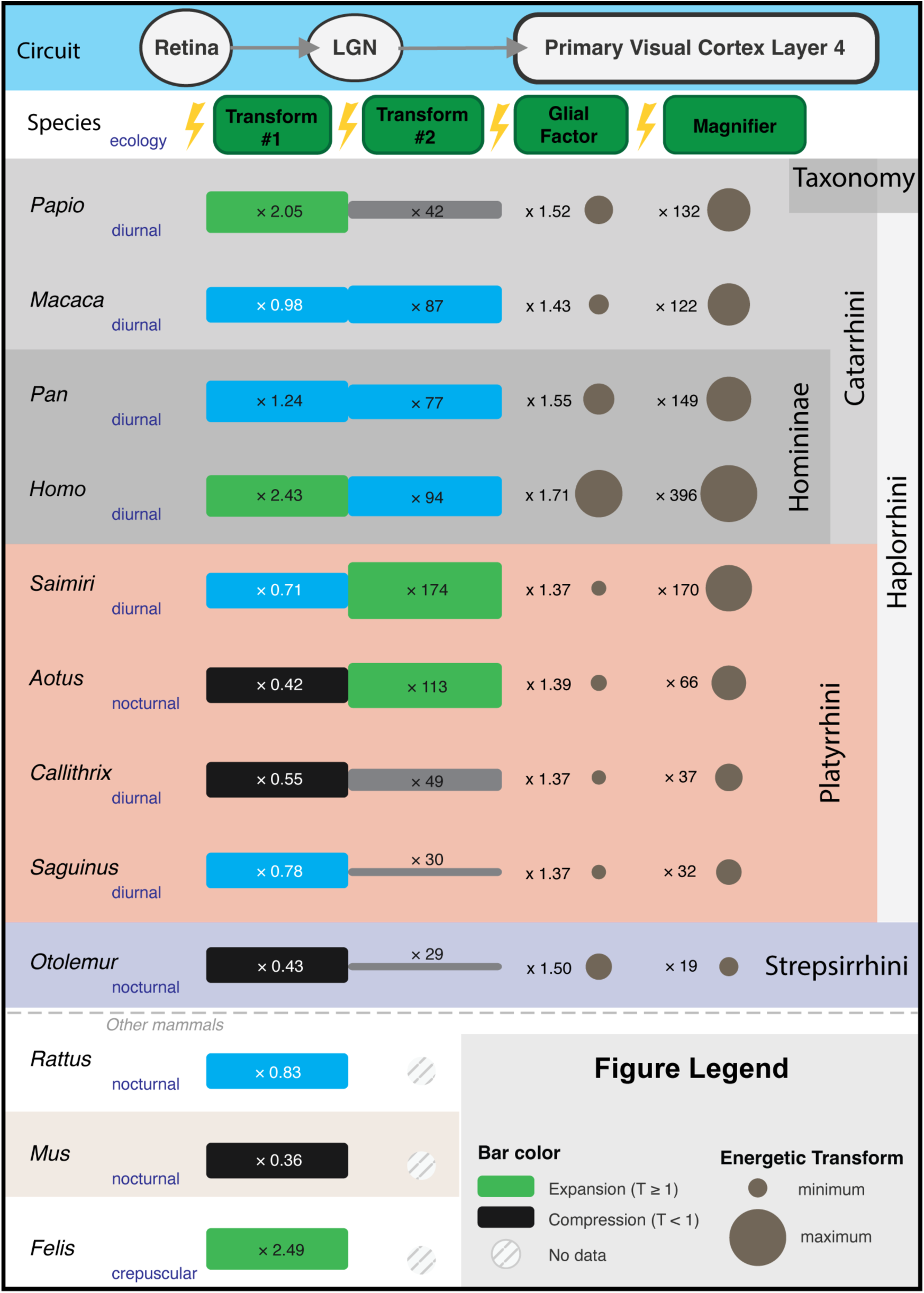
Ecological Energetics of the Primate Visual System. This schematic summarizes the stepwise expansion and compression of cell populations along the primary visual circuit. Specifically, we show circuit modifications from the retinal ganglion cells (RGC) through the lateral geniculate nucleus (LGN) to V1 layer 4 for all nine primates and three mammals for comparison. The top banner depicts the circuit topology (Retina → LGN → V1 layer 4), with lightning-bolt symbols at each relay denoting the energetic cost of maintaining the local cell population. Species are arranged phylogenetically within background-shaded bands: Catarrhini (grey; *Papio*, *Macaca*) including Homininae (*Pan*, *Homo*), Platyrrhini (salmon; *Saimiri*, *Aotus*, *Callithrix*, *Saguinus*), and Strepsirrhini (blue-grey; *Otolemur*). Activity pattern (diurnal, nocturnal, or crepuscular) is indicated beneath each genus name. Three non-primate species (*Rattus*, *Mus*, *Felis*) are shown below the dashed line for comparison, with hatched bars indicating that LGN-to-cortex data are unavailable for these taxa. Four quantities are displayed for each species. Transform #1 depicts the retina-to-LGN cell ratio as a horizontal bar whose color indicates whether the transform represents an expansion (green, T1 well above 1), near equivalency (blue, T1 ∼ 1) or a compression (black, T1 well below 1). Transform #2 depicts the LGN-to-layer 4 neuron ratio as a second bar with the same color scale. The GNR column shows the glial amplification factor, defined as (1 + GNR), displayed both as a numerical multiplier and as a scaled filled circle whose area is proportional to the value, and captures the additional metabolic burden enabled by the glial population associated with each neuron. The magnifier term is computed as T1 × T2 × GNR and displayed at the far right as a scaled filled circle representing the total signal amplification predicted for V1 layer 4 neurons from the periphery.

Retinal energetic regime predicts V1 neuron density across the 15-species sample (Figure 4, Table 2, Table S1). In a pre-specified PGLS of log V1 neuron density on log daily retinal ATP expenditure and a primate/non-primate binary variable (A3), daily retinal ATP was negatively associated with neuron density (slope = −0.386, SE = 0.112, p = 0.005), and primates showed a systematically higher neuron density than non-primates at equivalent retinal expenditure levels (intercept shift = +0.565, SE = 0.070, p = 3.48 × 10⁻⁶). Together, the two predictors account for 85% of interspecific variance in V1 neuron density (PGLS R² = 0.849, Pagel’s λ < 0.001). This result was stable in sign and significance across 100 posterior phylogenetic trees and across all jackknife deletions (Table S6).

**Figure 4.**
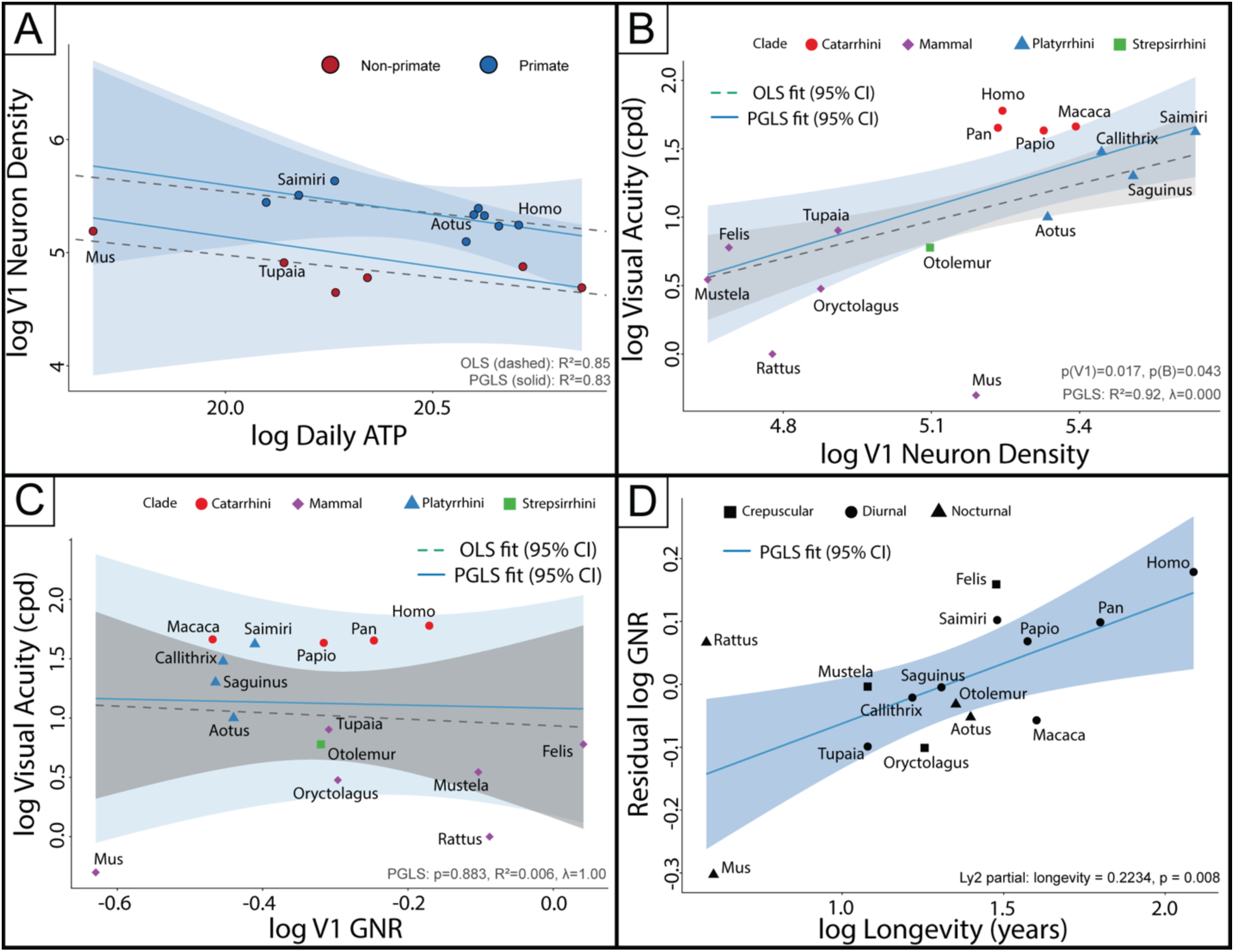
Axes of performance and maintenance in mammalian V1 cell composition. The depicted panels trace the relationships among retinal energy, cortical architecture, visual acuity, and lifespan across 15 mammalian species. Specifically, panels A through D together establish that V1 cellular composition decomposes into orthogonal axes: (A) an ecological axis linking retinal energy to cortical neuron density, (B) a performance axis linking neuron density to visual acuity, and (C, D) a maintenance axis along which glial investment tracks lifespan rather than visual resolution. Panel A shows how ecology shapes cortical architecture with plots of log V1 neuron density against log daily retinal ATP expenditure, with species coded as primate (blue) or non-primate (red). Panel B shows the performance axis with a plot of log visual acuity (cycles per degree) against log V1 neuron density, with brain mass controlled in the underlying PGLS model (Dy4, Table 2). Points are colored by clade (catarhine, red; platyrhine, blue; strepsirhine, green; non-primates, purple), and both OLS (dashed) and PGLS (solid) fits are shown with 95% confidence intervals (shaded). Panel C shows the affordability null hypothesis with plots of log visual acuity against log V1 GNR with points colored by clade as in panel B. Panel D shows the maintenance axis with a plot of the residual log GNR (after controlling for V1 neuron density) against log maximum longevity (years) for all 15 species (Ly2, Table 2). Points are coded by activity pattern (circles, diurnal; triangles, nocturnal; squares, crepuscular). All models are phylogenetic generalized least squares with maximum-likelihood estimation of Pagel’s λ. Full model coefficients are reported in Table 2, with diagnostic plots and sensitivity analyses in Supplemental Blocks S3–S5.

GNR scales negatively with V1 neuron density across the same sample (model B4: PGLS slope = −0.490, SE = 0.143, p = 0.005, R² = 0.473, λ = 0.617, Table 2, Figures S9–S12, Table S1) such that taxa with the dense neuronal packing tend to have fewer glia. B4 is significant in 100 of 100 posterior phylogenetic trees with the negative slope stable across all trees. We tested whether V1 neuron density and GNR each predict behavioral visual acuity using PGLS models with log-transformed variables (Figure 4, Table 2, Supplemental Table S4). In a model regressing log visual acuity on log V1 neuron density and log brain mass (model Dy4), V1 neuron density was a significant positive predictor of acuity (partial slope = 0.806, SE = 0.327, p = 0.032), as was brain mass (partial slope = 0.423, SE = 0.099, p = 0.001). Brain mass was included to account for allometric size variation across species, and together the two predictors account for 65% of interspecific variance in visual acuity (PGLS R² = 0.651, Pagel’s λ = 0.742; Table 2). In leave-one-out jackknife analyses, the neuron-density partial coefficient remained significant in 11 of 15 deletions, where four deletions (*Aotus*, *Saimiri*, *Rattus*, *Oryctolagus*) moved it to non-significance, but the sign never changed (Table S6).

In a model regressing log visual acuity on log GNR alone (model Dy1), GNR was not a significant predictor (slope = 0.230, SE = 0.181, p = 0.221, R² = 0.113, λ = 1.000; Table 2). GNR remained non-significant when tested alongside brain mass (Dy2) and alongside both neuron density and brain mass (Dy3; Table S3). The acuity models therefore indicate that neuron density but not GNR carries predictive information about visual resolution, whereas GNR does not. We tested whether GNR is associated with species maximum longevity after accounting for the structural GNR–neuron density relationship established by B4. Maximum longevity was selected as the life-history variable most directly relevant to the cumulative duration of circuit operation (24–27,30,40–42,44), while recognizing it is an indirect measure.

In a PGLS (Ly2) of log GNR on log V1 neuron density and log maximum longevity jointly (Figure 4, Table 2), both predictors were highly significant. V1 neuron density was negatively associated with GNR (partial slope = −0.510, SE = 0.073, p = 1.40 × 10⁻⁵), consistent with B4. Maximum longevity was positively associated with GNR after controlling for neuron density (partial slope = 0.262, SE = 0.056, p = 5.57 × 10⁻⁴). Together, the two predictors account for 82% of interspecific variance in GNR (PGLS R² = 0.818, Pagel’s λ < 0.001). The longevity association was replicated within the primate subsample (model Ly7: slope = 0.321, SE = 0.088, p = 0.008, R² = 0.656, λ < 0.001, N = 9, Table 2). The within-primate signal is illustrated by humans (*Homo*) and our sister taxa (*Pan*), which have nearly identical V1 neuron densities (∼175,000 neurons/mm³) but diverge in maximum lifespan and GNR (∼0.7 vs. ∼0.566). Both predictors in Ly2 retained significance under all 15 leave-one-out species deletions, including deletion of humans (neuron density maximum p = 1.53 × 10⁻⁴; longevity maximum p = 0.009; Table S6), and across 100 posterior phylogenetic trees (Table S5). Adding longevity to the acuity model (Ly6) yields no improvement (p = 0.959; Table S8), confirming that the two associations are empirically distinct. We report a secondary model testing lifetime retinal ATP expenditure (Table S8).

All principal findings were robust across 100 posterior phylogenetic trees, leave-one-out jackknife resampling, ±20% energy scenario perturbations, fixed-λ sensitivity analysis, and Cook’s distance diagnostics (full details Tables S4-S8). Humans have a V1 GNR substantially above the B4 prediction for its moderate neuron density, and the Ly2 model does not fully account for the magnitude of this departure, suggesting a residual elevation beyond what neuron density and longevity together predict. Independent studies in human V1 (37–39) and frontal cortex (30) are consistent with the elevated GNR reported here.

## Discussion

Across 15 mammalian species spanning five orders, V1 cellular composition varies along two partially separable dimensions. V1 neuron density, together with brain mass, predicts behavioral visual acuity (Dy4: R² = 0.651, density partial p = 0.032), whereas the glia-to-neuron ratio shows no reliable relationship to acuity in the same phylogenetic framework (Dy1: p = 0.221). By contrast, the glia-to-neuron ratio covaries positively with species maximum longevity after accounting for neuron density (Ly2: R² = 0.818, longevity partial p = 5.57 × 10⁻⁴), a pattern that remains significant within primates alone (Ly7: N = 9, p = 0.008) and is robust across 100 posterior phylogenetic trees and all 15 jackknife deletions. Adding longevity to the acuity model does not improve fit (Ly6: p = 0.959), indicating that the acuity and longevity associations are empirically distinct rather than secondary reflections of a shared size or phylogenetic gradient. Together, these results indicate that the cellular features predicting visual performance are not the same as those predicting the long-term maintenance of that function.

The acuity result is consistent with, and extends, predictions from the cortical sampling literature. Stevens (12) showed that V1 neuron number scales geometrically with LGN input size across haplorhines as a consequence of mapping a two-dimensional retinal surface onto a three-dimensional cortex, but he did not test whether this variation predicts what an animal can actually resolve. Srinivasan, Carlo, and Stevens (13) supplied the missing functional prediction because under by Nyquist-sampling logic, denser neuronal packing should support finer spatial resolution. Dy4 confirms that prediction across a five-order mammalian sample. In Dy4, brain mass is not a nuisance covariate but the term that captures shared allometric variance with body size, allowing neuron density to emerge as the functionally relevant component. The residual phylogenetic signal (λ = 0.742) further indicates that acuity also reflects inherited clade-level factors, including optical quality, foveal specialization, and post-V1 circuitry. Consistent with that interpretation, model A3 shows that daily retinal ATP expenditure and clade membership together predict 85% of the variance in V1 neuron density (R² = 0.849, p = 0.005), placing the cortical acuity axis within an ecological chain from photoreceptor complement through retinal energetics to cortical cellular organization and behavioral choice (11–13,21,23,29,31,32,45).

In contrast to neuron density, GNR shows no statistical relationship to visual acuity. Within the same phylogenetic framework, species sample, and response variable that recover the neuron-density effect in Dy4, Dy1 detects no GNR–acuity association (p = 0.221), and GNR remains non-significant after controlling for brain mass (Dy2: p = 0.615). This asymmetry argues against a generic power limitation, which should weaken inference for both cortical predictors rather than selectively eliminate the glial signal. Instead, the Dy1 null is consistent with the retinal dissociation formalized by Heras and Laughlin (11), in which spatial performance parameters are largely independent of the energetic cost of maintaining the photoreceptor array. The present data extend that logic to cortex: V1 neuron density aligns with the performance dimension, whereas GNR does not. Model B4 then identifies what GNR does track, showing a negative scaling relationship with V1 neuron density across the 15-species sample (slope = −0.490, SE = 0.143, p = 0.005, R² = 0.473), consistent with the cross-structure scaling law reported by Mota and Herculano-Houzel (34,45). Species with the densest neuronal packing therefore maintain fewer glia per neuron, suggesting lower per-neuron support demands under compact cellular conditions. Against this structural baseline, the longevity analysis asks whether residual GNR variation carries an additional life-history signal.

After controlling for the negative GNR–neuron density relationship established by B4, GNR is a strong positive predictor of species maximum longevity (Ly2: longevity partial slope = 0.262, SE = 0.056, p = 5.57 × 10⁻⁴; full model R² = 0.818, λ < 0.001). This coefficient is a robustly recovered parameter in the study, retaining both sign and significance across all 15 jackknife deletions (maximum p = 0.009) and across 100 posterior phylogenetic trees. Importantly, this result is modulated by brain mass which also associates with longevity. The association is also detectable within primates alone (Ly7: slope = 0.321, p = 0.008, N = 9), indicating that the cross-order result is not driven solely by rodent–haplorhine contrast. Together, these findings support the interpretation that glial allocation reflects the cumulative metabolic burden of sustaining cortical function across the lifespan. That interpretation is biologically plausible because astrocytes, oligodendrocytes, and microglia each contribute to forms of neural maintenance whose costs accumulate over time rather than at a single moment of performance (11,27,28,31,44,46). Consistent with this view, Sherwood et al. (30) and Pelvig et al. (47) reported elevated glial fractions in human cortex relative to allometric expectation, a pattern the present V1 data extend to a defined sensory area with direct stereological measurement. Still, maximum longevity is only an indirect life-history proxy, so these comparative data support a maintenance-related interpretation without resolving the cellular mechanism. More direct tests will require cell-type-resolved counts across species paired with metabolic rate measurements.

Among the nine primate genera, *Homo sapiens* has the highest GNR in the sample (V1 mantle: 0.675; layer 4: 0.734), well above the nonhuman primate range (mantle: 0.340–0.566; layer 4: 0.366–0.553). This elevation is driven entirely by glial excess, not neuronal depletion: human V1 glia density is the highest among primates (118,336 glia/mm³), whereas human neuron density (175,332 neurons/mm³) is nearly identical to that of *Pan troglodytes* (171,638 neurons/mm³; ratio = 1.02). The human outlier therefore requires a process that selectively elevates glial abundance beyond what local neuronal workload alone would predict. Ly2 explains part of this departure, because the positive longevity term shifts predicted human GNR above the primate centroid, but *Homo* remains detectably above the regression surface even after longevity is controlled. Consistency with independent stereological reports from human V1 (37,38) and frontal cortex (30) argues against simple measurement artifact. The macaque data of Giannaris and Rosene (48) provide only partial reassurance on that point. The current evidence therefore indicates that humans combine moderate V1 neuron density and moderate primate acuity with the highest glial support per neuron in the sample, a pattern more consistent with long-term maintenance demands than with spatial resolution.

Several limitations qualify these conclusions. Although the 15-species, five-order sample is sufficient to recover large and phylogenetically robust effects, the neuron-density partial in the acuity model loses significance when *Aotus*, *Saimiri*, *Rattus*, or *Oryctolagus* is removed individually, indicating that the performance-axis result depends on adequate sampling across the acuity range and would benefit from additional non-primate diurnal taxa. The human GNR estimate is derived from a single aged specimen, limiting inference about within-species developmental and aging trajectories. Because all analyses are restricted to V1, it also remains unknown whether the same dissociation holds in cortical areas with different functional demands or maintenance burdens. Finally, the retinal energy model assumes uniform per-neuron ATP costs across species; sensitivity analyses indicate that this simplification does not qualitatively change A3, but species-specific metabolic data will ultimately be required.

The central implication of these findings is that mammalian V1 cellular composition does not vary as a single composite trait. Instead, across the species sampled here, V1 varies along at least two partially separable cellular axes of visual performance and the maintenance of function. The constructional basis for this dissociation can be found in the alternative neurogenic and gliogenic developmental transcriptional programs that pattern cortex through partially separable temporal cascades (41,42,49). Whether the same two-axis organization extends to other cortical areas, including V2, MT, and prefrontal cortex, each with distinct functional demands and species-specific lifespan exposure, remains an open empirical question. Future work combining lifespan-spanning human V1 samples, cell-type-resolved stereology that distinguishes glial subtypes, and species-specific metabolic rate measurements would provide more direct tests of the maintenance interpretation and a firmer account of the residual human elevation.

## Materials and Methods

This was an archival study and used no live animals. Stereological data on the cellular composition of primary visual cortex (V1) was collected by DJM from nine primate genera (Table 1) using the StereoInvestigator system (MBF Bioscience) on a Nikon E80i microscope with a motorized stage (50). Specimens were selected from the Kaas laboratory collection only when the full extent of V1 was accessible in at least one tissue series stained for Nissl, and we obtained regional volumetrics and cell counts by type as described (30,35,43). Briefly, thalamorecipient layer 4 was defined following Hassler’s (4) classification as the parvorecipient (4b) and magnorecipient (4a) subdivisions (1,2) that corresponds to known (3,46,51) molecular patterns (Figures S1-S4) that surround the small round neurons that classically (1) defined the granular layer (Figures S5-S8). The glia-to-neuron ratio (GNR) was computed for each region as total glia count divided by total neuron count. Cell densities (neurons or glia/mm³) were computed by dividing total counts by the Cavalieri volume estimate. DJM collected all estimates under design, with a coefficient of error CE < 0.1 (Gundersen m = 1) (35,52).

The primary stereology data were then contextualized by surveying the literature (16,37–39,48,53) on the visual system to construct a 15 taxa dataset (Table 1) to test the first principles hypotheses of V1 cellular composition that also emerged (Table S1) from our survey of the literature (8,11–16,20,21,23). Specifically, visual acuity values (cycles per degree, behavioral threshold) were compiled from Veilleux and Kirk (54), cross-referenced with Prusky et al. (55) for rodents. Maximum longevity was obtained from the AnAge database (24,56). Body mass, brain mass, eye axial length, and temporal niche were compiled from various published sources, and full source documentation is available in supplemental information (see Supplemental Methods).

Daily retinal ATP expenditure (Edaily) was computed by summing photoreceptor maintenance costs across illumination phases (scotopic, mesopic, photopic) for a circadian cycle or 24 hr period. Energetics estimates (11,21,23,29) were fully derived from the literature for rods (22) and cones (57) along with comparative estimates of counts where available and anatomical organization for phylogenetic context (8,16–18). Phase durations were estimated from published sleep durations and temporal-niche classifications (Table S3). This variable condenses photoreceptor complement, retinal size, and activity pattern into a single index of peripheral metabolic demand rather than an estimate of absolute expenditure. Sensitivity to ±20% perturbations of phase-duration assignments is reported in Supplemental Table S7.

Species relationships were represented by the maximum clade credibility consensus tree from the posterior distribution of Upham et al. (58), pruned to the 15 study species (4,098 species, root age 78.3 Ma). All continuous variables were log₁₀-transformed, and models were fit under both ordinary least squares (OLS) and phylogenetic generalized least squares (PGLS) using R packages ape, caper, and geiger (59,60). Pagel’s λ was estimated by maximum likelihood (61) and Blomberg’s K computed as a complementary phylogenetic signal statistic (62). Full details are in supplemental information (Tables S4–S7).

Statistical tests of principled hypotheses (63) were conducted to assess three queries: first to test relationships to ecology, second to test relationships to brain size, and third to test for relationships to life history. Model comparisons used AICc (64), and six classes of sensitivity analysis were conducted, using fixed-λ PGLS and Cook’s diagnostic, with full details available in the supplemental information.

### Sample

The complete species-level dataset for V1 neuron and glia densities, retinal and LGN cell counts, and ecological variables, is presented in Table 1 with primary model results in Table 2. Our stereological study sample comprised original stereological data on the cellular composition of primary visual cortex (V1) in nine primate genera. Sample sizes are uneven across taxa, reflecting the availability of specimens in which the entire extent of V1 was present for study with at least one series of sections stained with Nissl’s method in the Kaas collection. No live animals were used in this work as this was an archival study of previously collected tissue accompanied by a literature review. All procedures involving animals and animal tissues that ultimately contributed histological material imaged during this project followed the NIH Guidelines for the Use of Laboratory Animals and were approved in advance by the Vanderbilt University Institutional Animal Care and Use Committee when specimens were not received from outside institutions (35,43,65). The human specimen was obtained from an 82-year-old female through the Anatomical Request Program at the University of Tennessee Memphis Health Sciences Center (Memphis, TN, USA). The baboon and chimpanzee specimens were purchased from the Texas Biomedical Research Institute’s Southwest National Primate Research Center (San Antonio, TX, USA). All other specimens were archival from our group’s previous experiments (35,65).

We calculated the phylogenetic neuroanatomical context for V1 cell count data from the literature (Figures 2 & 3; Table S1). Specifically, the evolutionary biological context for the V1 data collected is the comparative organization of the visual system, in particular the canonical path from retina to lateral geniculate nucleus in the thalamus (LGN) to primary visual cortex layer 4 (3,16). We measured the transform of signal from the retina to LGN as T1, and the transform from the LGN to V1 layer 4 as T2 (Figure 3, Table S2). Together with the primary stereology work, this literature enriches our comparative cell counts dataset for a more informative phylogenetic analysis. Before testing the main predictions, we first characterize the distribution of V1 cellular variables and visual acuity across this full sample and report their phylogenetic signal to establish the comparative landscape within which the model tests are situated.

### Histology

All specimens had, at minimum, a single series of sections stained with thionin for Nissl substance, on which the majority of stereological counts were collected. Neighboring section series stained for cytochrome oxidase (CO), acetylcholinesterase (AChE), neuronal nuclei (NeuN; AB667, Millipore, Billerica, MA, USA), parvalbumin (PV; P3088, Sigma-Aldrich), and vesicular glutamate transporter 2 (VGLUT2; Millipore) were not always available across the entire sample but were used when available to identify layer 4 following Hassler’s classification (1,2,4). Where available, molecular markers facilitated laminar delineation more precisely than Nissl staining alone (46,51,66,67), and the use of multiple histological preparations together with previously published descriptions enabled the identification of V1 and granular layer 4 in each primate brain (4,8,39,68).

We defined V1 by its highly laminated structure, in which the outer line of Baillarger—the stria of Gennari—is evident even prior to histological processing (1,4,69–71), and which can be further identified by the characteristically dense staining of CO, AChE, PV, and VGLUT2 across the cortical mantle (2,35). The cortical mantle was defined as the tissue between the pial surface and the transition to white matter (WM), a boundary identifiable across histological preparations as the point at which the neuron-rich gray matter gives way to the axon-rich white matter (Supplemental Figures S1-S4). We defined the granular cell layer 4 (L4) in the traditional manner (1,2,4,37,66) by the high density of small, round neurons visible in preparations for Nissl and NeuN (Supplemental Figures S5-S8), supplemented by staining for VGLUT2 and PV, which label terminals and neurons heavily innervated by the LGN (2,38,68,72–74), and for CO and AChE, which reveal more diffuse synaptic input (46,50,51,75).Throughout this study, layer 4 refers to Hassler’s (4) definition: the parvorecipient (4B) and magnorecipient (4A) subdivisions of the granular tier in layer 4, exclusive of Brodmann’s 4A and 4B, which correspond to layer 3 in Hassler’s schema (2,8).

Thionin staining for Nissl substance revealed cellular morphology used to distinguish neurons from glia and constituted the basis for the majority of counting in this study (30,35,37). Immunolabeling with NeuN selectively visualized neurons (67,76) and was also used for optical fractionator counts in specimens where NeuN-stained series were available (35,52,77). In Nissl-stained material, neurons were distinguished from glia by the presence of a darkly stained nucleolus within a more moderately stained nucleus, surrounded by a paler-staining perikaryon that was typically, but not always, larger than other cellular profiles in the field of view (30,37,38,78). Glial cells (astrocytes and oligodendrocytes) were identified by the absence of a visible nucleolus, a uniformly stained nucleus, and the absence of appreciable cytoplasm surrounding the nucleus, while microglia, pericytes, and endothelial cells were not included in the analysis.

### Stereology

All measurements and images were collected using a computerized stereology system consisting of a Nikon E80i microscope outfitted with ×2 to ×100 oil-immersion (NA = 1.3) objective lenses, connected to a motorized stage (Ludl Instruments) running StereoInvestigator software (MBF Bioscience, Williston, VT) on a Dell desktop workstation (35,50). Regional volumes were estimated using the Cavalieri method, and cell numbers were estimated using the optical fractionator with StereoInvestigator. Contours delineating V1 and layer 4 were drawn at ×2 magnification, and all counting and high-resolution image acquisition were performed at ×100 oil immersion under Köhler illumination. A single investigator (DJM) performed initial pilot tests in each species to establish stereological sampling parameters that maintained a low coefficient of error (Gundersen m = 1 CE < 0.05), as before (35,52). Intraclass correlation coefficients (ICC) were assessed following published guidelines (79), and DJM achieved an intra-rater reliability of 0.92 (n = 7, 95% CI: 0.88–0.96), indicating good agreement per established benchmarks (52). The final parameters were as follows: number of sections investigated per specimen, 9–12 (mode = 10); counting frame area, 15–30 µm² (mode = 15 µm²); optical dissector height, 6–10 µm (mode = 6 µm); guard zone, 3 µm; frame step for V1 mantle, 500–2,000 µm²; frame step for layer 4, 200–350 µm²; and frequency of mounted section thickness measurements, 1 in ∼8. Cavalieri point-counting grid spacing ranged from 200 µm² in the smallest brains to 1,000 µm² in the largest (human, chimpanzee). More than 300 probes per region of interest have been shown to reduce the variance of estimates even when the underlying distribution is non-normal to maintain data quality. Accordingly, a mean of 311 optical dissector probes (SEM = 39) were investigated per specimen, and the top of the cell soma was used as the unique identifying feature of each object to avoid double-counting (30,35,36). Total cell numbers were estimated by multiplying the number of counted cells by the reciprocals of the area section fraction, the serial section fraction, and the tissue section fraction (52).

### Derived cellular variables

We calculated multiple derived measures from the primary datasets, including density measures, volumetric ratios, and cell type relationships. The volumetric proportion of layer 4 was computed as the ratio of layer 4 volume to V1 mantle volume for each taxa. These derived measures are subject to the propagation of variance from both numerator and denominator, but the high precision of the underlying counts constrains propagated error (CE ∼ 0.1). Cell densities (neurons/mm³ and glia/mm³) were computed by dividing the total cell counts by the corresponding Cavalieri volume estimate for each region and genus. The glia-to-neuron ratio (GNR) was computed for each region by dividing total glia count by total neuron count. The GNR provides a direct index of the metabolic infrastructure invested per unit of neuronal signaling capacity (28,30). Specifically, glia are required for neurotransmitter recycling, ionic homeostasis, and axon myelination (21,29,31,33,80–82). Accrodingly, the GNR captures some of the per-neuron overhead that the tissue must sustain during operation (28,30,80).

Following the tradition of a cortical magnification factor, we defined a magnifier index for the visual pathway (Table S2). Two circuit-level transform variables were computed to characterize signal amplification across the visual pathway to estimate a magnification index. Cell T1, the retinogeniculate transform, was defined as the ratio of LGN neuron count to retinal ganglion cell count, indexing whether the thalamic relay expands or compresses the retinal signal, and T1 values range from 0.36× in *Mus* (compression) to 2.49× in *Felis* (expansion). Cell T2, the geniculocortical transform factor, was defined as the ratio of V1 layer 4 neuron count to LGN neuron count, indexing the divergence of signal from thalamus to cortex, and T2 ranges from approximately 29× in *Otolemur* to approximately 174× in *Saimiri*, consistent with the scaling framework previously documented (12). Accordingly, we report on the phylogenetic variation of composite energetic cellular magnification at the level of the cerebral cortex to better understand how the retinogeniculate and geniculocortical relay transforms along with glial support reveal the total effective investment in V1 layer neurons per retinal ganglion cell (RGC) fiber. This magnification metric captures the total cellular investment—both neuronal and glial—per unit of peripheral input, weighted such that glia contribute proportionally to the metabolic load they impose on the neuronal substrate (28–30).

### Bioenergetic Framing

The analytical framework of this study treats the mammalian visual system as a network of energy-transducing elements whose measurable anatomy — cell counts, cell-type ratios, relay architecture — constitutes an inventory of the components through which energy flows during visual processing. This approach draws on the bond-graph tradition in bioengineering (83–86), which models biological systems as networks of energy storage, dissipation, and transduction connected by power bonds that enforce thermodynamic conservation across mechanical, electrical, and biochemical domains. We do not implement a formal bond-graph model, but we do adopt the thermodynamic conservation of energy that forms the organizing principle of biochemical reactions (87). Specifically, we propose the cellular composition of neural tissue is an empirical parameterization of the energy-flow network that produces neural function, and that cross-species variation in composition reflects how evolution has tuned the parameters of that network under ecological constraint (88). Specifically, biological systems can be described mathematically as consisting of elements that store and dissipate the flow of energy, and we adopt the view that any biosynthetic process (the molecular signaling cascades that drive Hebbian synaptic plasticity) contributes to an organism’s biomass. Critically, this formalism of energetics applies to neural signaling (21,29,82) and can be understood to supplant the mathematical relationships that are currently presumed to drive sensation and perception from a phenomenological standpoint (7,89–91), with a mechanistic parameterization (83,87,88). The proposed energetic framing of the mammalian visual system thus enables a first principles account of the relationship between ontogenetic biosynthetic events (41,42,88), like neurons of a particular flavor (92–97), and their receptive field properties (5–7,98) over evolutionary scaling (8,12,14,18,19,30,80,99).

An energetics perspective generates a specific cost decomposition for the visual pathway given the modern synthesis of evo-devo by molecular action (42,83,87,88,100,101). Any neural circuit imposes two categories of metabolic cost on the organism. First, each organism must grow and maintain the cellular neurobiological circuit, which is accounted for by the continuous expenditure of ATP required to maintain ionic gradients, drive synaptic transmission, and restore electrochemical homeostasis after each signaling event in a neuron (29,102–108). Attwell and Laughlin (21) quantified this cost at each relay of the visual pathway and showed that Na⁺/K⁺-ATPase–mediated gradient restoration dominates the total at every stage (29). The second is the maintenance cost of this signaling behavior, which is the cumulative burden of sustaining the cellular infrastructure — recycling neurotransmitters, maintaining myelin, regulating neurovascular coupling — across the organism’s lifetime (11,22,109). Dynamic energy budget (DEB) theory formalizes this distinction at the organismal level, partitioning total energy allocation into growth, operation, and maintenance, and predicting that maintenance costs accumulate as the temporal integral of the operating rate over the lifespan (110–114).

The neurobiology of this thermodynamic framing connects the two major classes of cells in the brain to alternative decompositions of metabolism. Specifically, the critical biological observation that connects energetics to measurable anatomy is the biosynthesis of functional cell types that have largely non-overlapping roles, as neurons differ from glia (41,42,81,115,116). Neurons are the substrate of perception and generate action potentials to transmit the synaptic signals that determine spatial organization and system operation, or biological behavior (115). Glial cells are historically known for metabolic support (28,30), the details of which turn out to constitutively modulate neuronal firing probabilities (32) by recycling glutamate, buffering extracellular ions, growing myelin (44), and coupling neural activity to vascular supply (11,31), completing the neurovascular unit (NVU) (117). If operating cost is driven primarily by neuron number (12,28,29), and maintenance cost is borne primarily by glia (80), then the two cell populations should index different dimensions of the total energetic accounting of perception (105,110,114,118). For instance, neuron density should index the instantaneous computational capacity of V1 — and therefore predict visual performance. The glia-to-neuron ratio should index the per-neuron maintenance overhead — and therefore predict not how well the animal sees but the capacity for ecological sensory challenge, or how the animal sees across ecological conditions. The thermodynamic framing of bond graphs thus provide a first-principles explanation of biological system operation as metabolic energy flows that map onto known cell classes.

Recent work has demonstrated an energetic dissociation of the neurobiology of the visual system. Specifically, Heras and Laughlin (11) demonstrated the disambiguation of performance from cost parameters in the retina. It turns out the optical dimensions that set spatial resolution are insensitive to the photoreceptor energy ‘tariff’ the authors propose, whereas the metabolically sustainable photoreceptor array depth is quite sensitive. The model structure of the present study tests whether the same dissociation characterizes the cortical field that processes the retina’s (via thalamus) output. The performance models (Dy series) test whether V1 neuron density predicts visual acuity while GNR does not. The maintenance models (Ly series) test whether GNR, after controlling for neuron density, tracks species longevity. The ecology-to-architecture model (A3) tests whether the peripheral energy regime — the daily metabolic demand imposed by the retina’s photoreceptor complement and illumination environment — predicts the neuron density of the cortex that receives its signal. Together, these models constitute a decomposition of V1 cellular composition into a performance axis (neuron density → acuity) and a durability axis (GNR → lifespan), with ecology acting on the system through the retinal energy budget.

The bioenergetic estimates reported in this study operate at the level of cells. Specifically, we used published per-event energy values (11,21,29,82,119) as rate constants rather than deriving them from thermodynamic first principles to model cost per action potential, cost per synaptic transmission, and cost per gradient restoration cycle. We use the term "mesoscale” for our energy accounting to distinguish this approach from the formal bond-graph methodology that operates at the level of individual molecular species and mathematical precision (83,84,87). The espoused mesoscale bond graph approach sacrifices molecular resolution for comparative applicability by permiting cross-species comparison of pathway cost topology using the same per-event constants, under the assumption that per-neuron signaling cost is approximately uniform across the taxa in our sample. This assumption is a necessary simplification and is evaluated in the sensitivity analyses and discussed as a limitation of the current work. The resulting estimates reflect the topology of energetic variation across species — which relays dominate, which species invest most, how ecology maps onto cortical architecture — rather than precise absolute metabolic expenditure. This topological information is sufficient for the comparative tests that constitute the core of the study, because the hypotheses concern the relative scaling of performance and maintenance components, not the absolute joules consumed by any individual cortex.

### Comparative data

Given the variation in methodological provenance across the comparative dataset, we took an inclusive approach to interpreting and adapting reports from the literature and tested for signal (Table S4). Primary data from DJM on V1 whole-mantle and layer 4 stereological estimates for nine primate genera were obtained by design-based stereology (Cavalieri volumetry and optical fractionator cell counts; CE < 0.05 for all estimates). Data from the literature also contained significant stereological estimates that we adopted directly (16,17), with minimal modifications given how the data were collected to ensure commensurability across studies. The least concordant estimates come from non-stereological methods that we treat here as previously (16), adjusting where overlap exists to generate ballpark estimates across as wide a sampling as possible (35,36). Only the retinal data for *Pan* was estimated from existing statistical metrics to provide some estimate where no data existed in the literature.

### Retinal data

Retinal ganglion cell (RGC) counts and retinal areas for eight primate genera and three non-primate mammals were drawn primarily from Franco et al. (120) and Finlay et al. (17), who provide the most comprehensive compilations of primate retinal cell populations. Species-specific rod and cone photoreceptor counts, which are required for the retinal energy model described below, were obtained from the same compilations, supplemented by Peichl (121) for non-primate taxa and Müller and Peichl (122) for the cone-dominated retina of *Tupaia belangeri*. The *Pan troglodytes* RGC count was estimated by PGLS regression from retinal area, as no published count exists for this genus.

### Lateral geniculate nucleus

Estimates of the visual thalamus or lateral geniculate nucleus (LGN) neuron counts were drawn primarily from Finlay et al. (16) and sources cited therein. The *Otolemur* LGN values were derived from Collins et al. (99) as also reviewed by Finlay et al. (16), with volume and count correction factors calibrated by overlapping species for which both fractionator and stereological estimates exist.

### V1 data

V1 cell densities for the six non-primate species in the expanded sample were obtained from publications reviewed. Stereological studies exist for *Mus musculus* neuron and glia densities from Keller et al. (123), who compiled mouse V1 cell counts from design-based stereological and isotropic fractionator studies (neuron density: 155,000/mm³; glia density: 36,364/mm³). Work in *Rattus* (neuron density: 60,020/mm³; glia density: 49,040/mm³) comes from with Gabbott and Stewart (124), with GNR cross-referenced to Yates et al. (53) and Juraska (125). Work in the cat (*Felis catus*) by Beaulieu and Colonnier (126) and Gabbott and Somogyi (127), who reported values in the range 48,000–54,210/mm³ for the binocular representation of area 17, from which we adopted 49,000/mm³ as the representative value. Cell composition estimates for *Mustela putorius furo* is taken from López-Hidalgo et al. (128), who reported a neuron-to-astrocyte ratio of 2.48 (astrocyte density: 17,896/mm³). For *Tupaia belangeri* and *Oryctolagus cuniculus*, V1 volume density estimates were derived from the approximately constant relationship between neuron number per unit cortical surface area and cortical thickness following the Stevens (13) tradition. Specifically, Srinivasan, Carlo, and Stevens (13) confirmed that non-primate mammals maintain approximately 97,780 neurons/mm² per unit of cortical surface in V1, a value consistent with the Rockel et al. (14) constant as replicated by Carlo and Stevens (129) using modern stereological methods. Specifically, division of this surface density by published cortical thickness values for each species yields estimated volume densities of approximately 81,500 neurons/mm³ for *Tupaia* and approximately 75,200 neurons/mm³ for *Oryctolagus*. Values from independent studies (37,38,48,130) were used for validation but not included in the primary dataset.

### Visual acuity

Published visual acuity values for each species (cycles per degree at behavioral threshold) were compiled from Veilleux and Kirk (54), who provided the most comprehensive compilations of behaviorally measured spatial acuity across mammals. For rodent species, acuity values were additionally cross-referenced with Prusky et al. (55) for *Mus* (0.5 cpd) and *Rattus* (1.0 cpd). All acuity values used in the analyses are behavioral thresholds (Table S1).

### Life history variables

Body mass, brain mass, and eye axial length were compiled from published sources for all 15 species (Table S1). Temporal niche (diurnal, nocturnal, or crepuscular) was assigned from published activity-pattern classifications. Maximum longevity values were obtained from the AnAge database (56), supplemented by species-specific literature where the AnAge entry was absent or contested. Sleep duration estimates were compiled from published sources for each species and activity budgets for two species (*Saguinus midas* and *Mustela putorius furo*) were estimated by analogy with closely related taxa for which published data were available (see Supplemental Table S1).

### Retinal energy model

The bioenergetic framework requires estimating the energetic demand that the retina places on the visual pathway. The retina operates continuously — photoreceptors maintain costly dark currents even during sleep — and thus, daily retinal ATP expenditure provides a summary index of the peripheral metabolic load that the cortex must process and sustain. Daily retinal ATP expenditure (Edaily) was computed for each species by summing the energetic cost of photoreceptor maintenance across four illumination phases: sleep, scotopic, mesopic, and photopic. Within each phase, energy consumption was calculated as the product of phase duration (in seconds), photoreceptor count, and the phase-appropriate ATP consumption rate. Rod ATP rates were taken from Okawa et al. (22) at 1.0 × 10⁸ ATP s⁻¹ rod⁻¹ in darkness, declining to 2.5 × 10⁷ ATP s⁻¹ rod⁻¹ under photopic illumination as the phototransduction cascade is progressively suppressed. Cone ATP rates were taken from Ingram et al. (57) at 1.7 × 10⁸ ATP s⁻¹ cone⁻¹ in darkness, declining to 1.1 × 10⁸ ATP s⁻¹ cone⁻¹ under photopic illumination, reflecting the smaller absolute savings in cone dark current relative to rods.

The duration of each illumination phase was estimated for each species from published sleep durations and temporal-niche classifications (Supplemental Table S1). For diurnal species, waking hours were assigned predominantly to photopic conditions with a 1–2 hour mesopic transition at dawn and dusk. For nocturnal species, waking hours were assigned predominantly to scotopic conditions with a comparable mesopic buffer. For crepuscular species, a larger proportion of waking time was allocated to mesopic and scotopic phases. These phase assignments represent a first-order approximation, as no published study directly partitions the 24-hour light environment into scotopic, mesopic, and photopic durations for most species in our sample, and the true photic environment experienced by a free-ranging animal varies with canopy cover, lunar cycle, and weather etc.

The retinal energy model provides a ballpark energetic substrate of vision ecology. Specifically, the energy model serves to condense the combined effects of photoreceptor complement (rod-to-cone ratio), retinal size, and activity pattern into a single continuous variable (Edaily) that indexes the daily metabolic demand imposed by the visual environment. This variable is used as a predictor of V1 neuron density in model A3, testing the hypothesis that the peripheral energy regime predicts the cellular architecture of the cortex that processes its output. The model is not intended to estimate absolute metabolic expenditure, but to capture the topology of energetic variation across species robust to the ±20% perturbation of activity-phase durations tested in the sensitivity analysis (Supplemental Table S7).

### Cortical energy estimates

To quantify the energetic cost of visual processing at the cortical level, we compiled per-event bioenergetic cost estimates following the framework of Attwell and Laughlin (29,82,119), and extended this to the phylogenetic variation of the visual system (15–18,99,131). The visual system in mammals is recognized to exhibit a ‘convergence-then-divergence’ organization. Specifically, the retinal signal is compressed at the thalamic relay and then massively expanded at the geniculocortical relay and into cerebral cortex and V1. The cost categories include phototransduction and dark-current maintenance in photoreceptor outer segments, graded-potential signaling at retinal ribbon synapses, spike generation and axonal conduction in retinal ganglion cells, synaptic transmission at the retinogeniculate and geniculocortical relays, action potential generation in LGN relay neurons and V1 layer 4 granule cells, and the gradient restoration that follows each signaling event and dominates the total metabolic cost at every stage. Under the uniform-cost assumption, variation in total pathway cost across species is driven by neuron count and GNR. The resulting metabolic estimates reflect the topology of cost variation — which relays dominate, which species are expensive, and how ecology maps onto energetic investment — rather than the absolute magnitude of metabolic expenditure. We adopt a 100-ms stimulus epoch activating a single retinotopic column as the standard unit for cross-species comparison, following the convention established by Attwell and Laughlin (21). We provide per-event costs and full derivation parameters in the supplemental materials which contain the companion bioenergetics tables.

### Phylogenetic comparative methods

Brain mass was included to account for allometric size variation that would otherwise confound the relationship between neuron density and acuity across species differing substantially in body and brain size. Species relationships (Table S5) were represented by the maximum clade credibility (MCC) consensus tree from the DNA-only, tip-dated Bayesian posterior distribution of Upham et al. (58) (4,098 mammalian species; Dryad DOI: 10.5061/dryad.tb03d03). The full tree was pruned to the 15 species in our sample, retaining original branch lengths (root age 78.3 Ma). We evaluated phylogenetic signal and scaling relationships using a dual-model approach in which each dataset was analyzed under both ordinary least squares (OLS) and phylogenetic generalized least squares (PGLS) regression, following published guidelines for comparative studies (63). All continuous variables were evaluated both as raw values and following log₁₀ transformation. Log transformation linearizes allometric relationships, stabilizes variance, and is standard practice in comparative biology for cell count and volume data that span orders of magnitude (16,30,37,120,131).

For each pairwise comparison, we report both OLS and PGLS regression fits with 95% confidence intervals (Table S4), along with the summary statistics needed to evaluate phylogenetic signal: R², AIC for both models, Pagel’s λ, and the regression slope under each model. PGLS models were implemented using the R packages "ape", "caper" and "geiger" (59,60), with Pagel’s λ estimated by maximum likelihood as a continuous measure of phylogenetic signal in model residuals (61). Pagel’s λ ranges from 0 (no phylogenetic signal, where residuals are independent across species) to 1 (residuals covary in proportion to shared evolutionary history as described by a Brownian motion model). In addition, we computed Blomberg’s K (62) as a complementary measure of phylogenetic signal in individual traits. Blomberg’s K = 1 indicates signal consistent with Brownian motion, where K > 1 indicates stronger phylogenetic conservatism than expected, and K < 1 indicates weaker signal than expected.

### Model structure

The analysis was organized into pre-specified and exploratory model sets (Table S2), following recommendations for transparent reporting in phylogenetic comparative studies (63). The model structure reflects the bioenergetic framework described above, in which the visual pathway is decomposed into (1) how does the retinal energy regime relate to cortical composition, (2) how does cortical composition predict what the animal can see, and (3) how does cortical composition relate to biological operation timescales (i.e. ontogeny).

Three pre-specified model sets addressed the primary research questions. First, set A tested the ecology-to-organization mapping where V1 neuron density was the response variable to test whether cell number is predicted by brain mass, retinal energy expenditure, or a primate/non-primate binary variable. Second, set B tested the scaling relationship between glial investment and neuronal substrate where GNR is set as the response variable against brain mass or V1 neuron density. Third, in set C, the magnification factor is the response variable to test if a composite cellular investment metric scales with brain mass or some other predictor.

Additional models were parameterized to test for relationships between ecology and visual performance over the lifetime with cell composition. Ecological-load models (designated Ax, Bx, Cx) decomposed retinal energy expenditure into photopic fraction, ecological intensity, and dark-tax components to test whether the internal structure of the energy budget provides additional explanatory power beyond the aggregate. Additional visual-performance models (designated Ay, By, Cy, Dy) tested whether V1 cellular variables predict visual acuity. The maintenance models (designated Ly) tested the longevity prediction (Table S8). Each model set is clearly demarcated in all tables and supplemental materials, and model comparison within each set used the corrected Akaike Information Criterion (AICc) following Burnham and Anderson (64), with ΔAICc values computed relative to the best model within each response-variable set.

We conducted multiple classes of sensitivity analyses to assess the stability of results under alternative analytical conditions. First, all pre-specified models were refitted across 100 posterior trees (Table S5) drawn at random (seed = 12345) from the full 10,000-tree Bayesian posterior distribution of Upham et al. (58). Convergence was verified by comparison with a 200-tree sample, and coefficient signs, significance levels, and model rankings were stable across both samples.

Next, we did a leave-one-out jackknife analysis (Table S6) and removed each of the 15 species in turn, re-estimating model coefficients and significance under each deletion to identify influential taxa. Then we adjusted all energy-dependent models by refitting under ±20% perturbations of the activity-phase durations to test the sensitivity of the ecology–structure link (model A3) to uncertainty in the retinal energy estimates (Table S3). All PGLS models were re-estimated (132) under fixed values of λ (0, 0.5, 1.0) to confirm that results are not artifacts of the maximum-likelihood λ estimation procedure (Table S7).

## Acknowledgments

The authors thank Huixin Qi, Iwona Stepniewska, Anna Huang, Roshan Poudel, Pooja Balaram, Nicole Young, Jamie Reed, Chia-chi Liao, and Laura Trice for technical assistance. DJM acknowledges their wife and partner Dr. Zoe LeBlanc for without her integrity, love, and unwavering support this project would not have been possible.

## Supplemental Information

**Figure S1.**
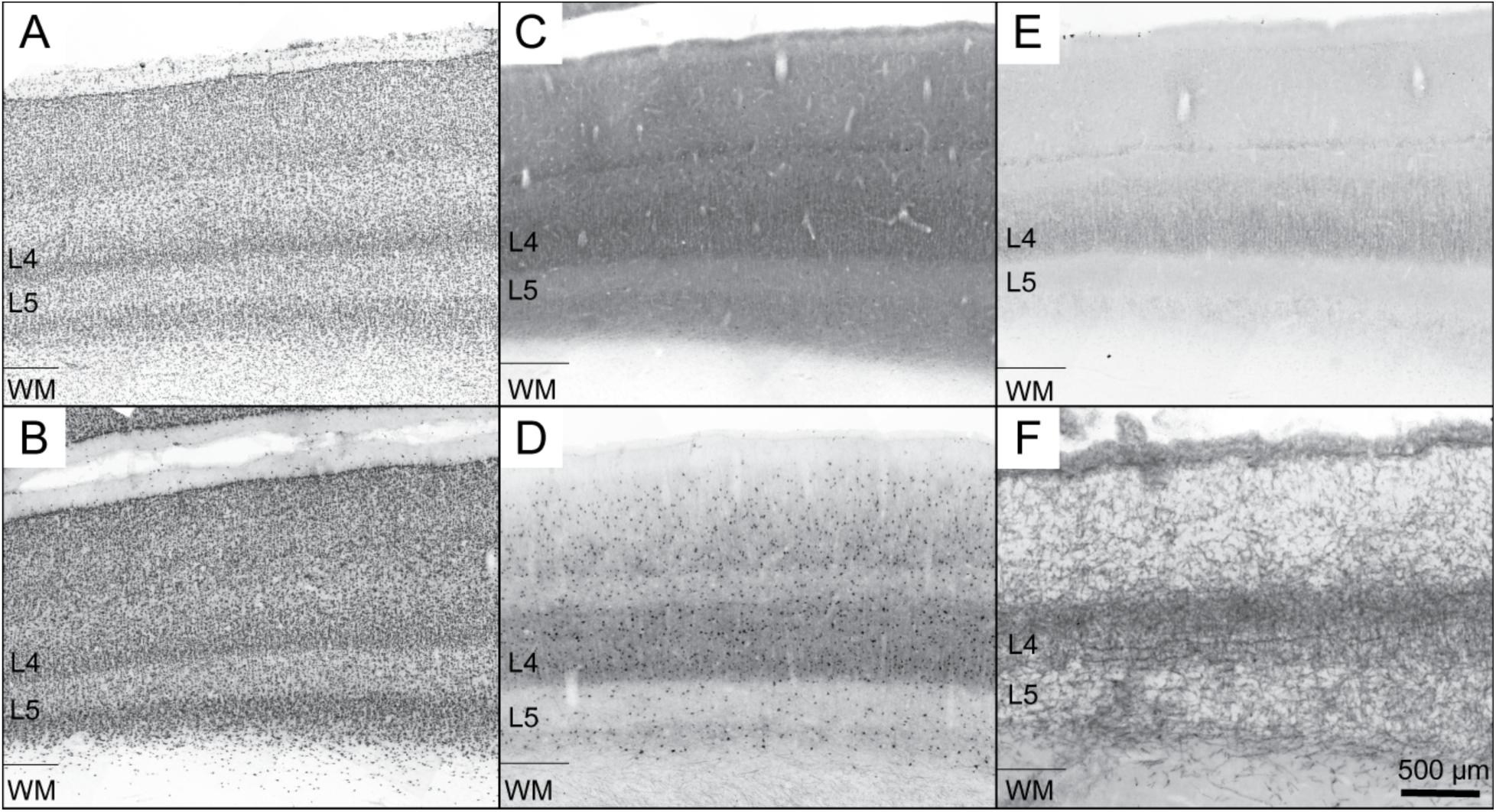
Characterization of layer 4 within V1 for *Macaca mulatta*. Low-magnification composite images (×10 objective) of adjacent coronal sections through V1 in the macaque monkey (*Macaca mulatta*), illustrating the laminar position and histological signature of the granular cell layer 4 (L4) as identified across six histological preparations. (A) Nissl (thionin); (B) neuronal nuclei antigen (NeuN); (C) cytochrome oxidase (CO); (D) parvalbumin (PV); (E) vesicular glutamate transporter 2 (VGLUT2); (F) acetylcholinesterase (AChE). L4, layer4, L5, layer 5, WM, white matter. Scale bar = 500 µm.

**Figure S2.**
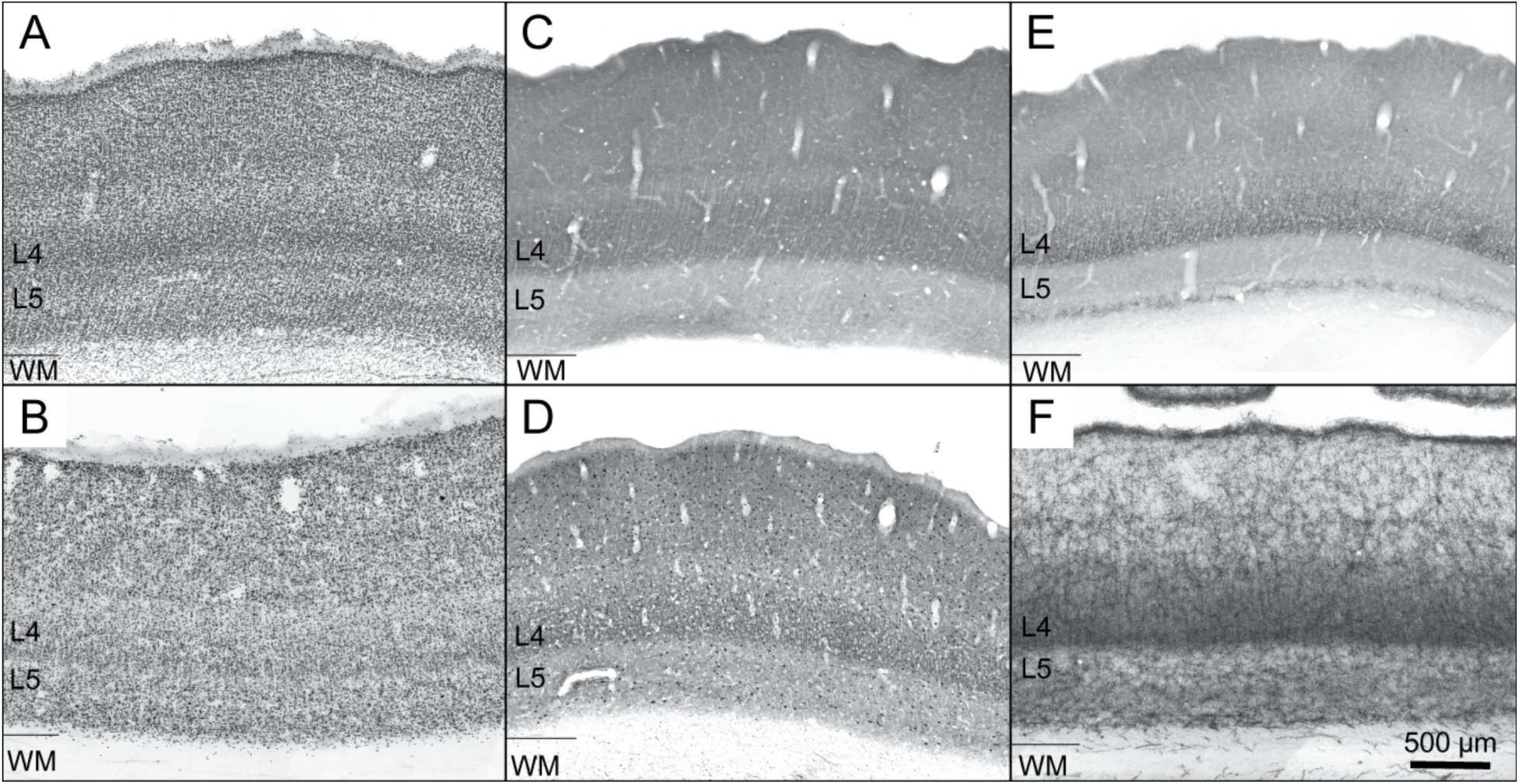
Characterization of layer 4 within V1 for *Saimiri sciureus*. Low-magnification composite images (×10 objective) of adjacent coronal sections through V1 in the squirrel monkey (*Saimiri sciureus*), illustrating the laminar position of the granular cell layer 4 (L4). (A) Nissl (thionin); (B) NeuN; (C) cytochrome oxidase; (D) parvalbumin; (E) VGLUT2; (F) acetylcholinesterase. L4, layer4, L5, layer 5, WM, white matter. Scale bar = 500 µm.

**Figure S3.**
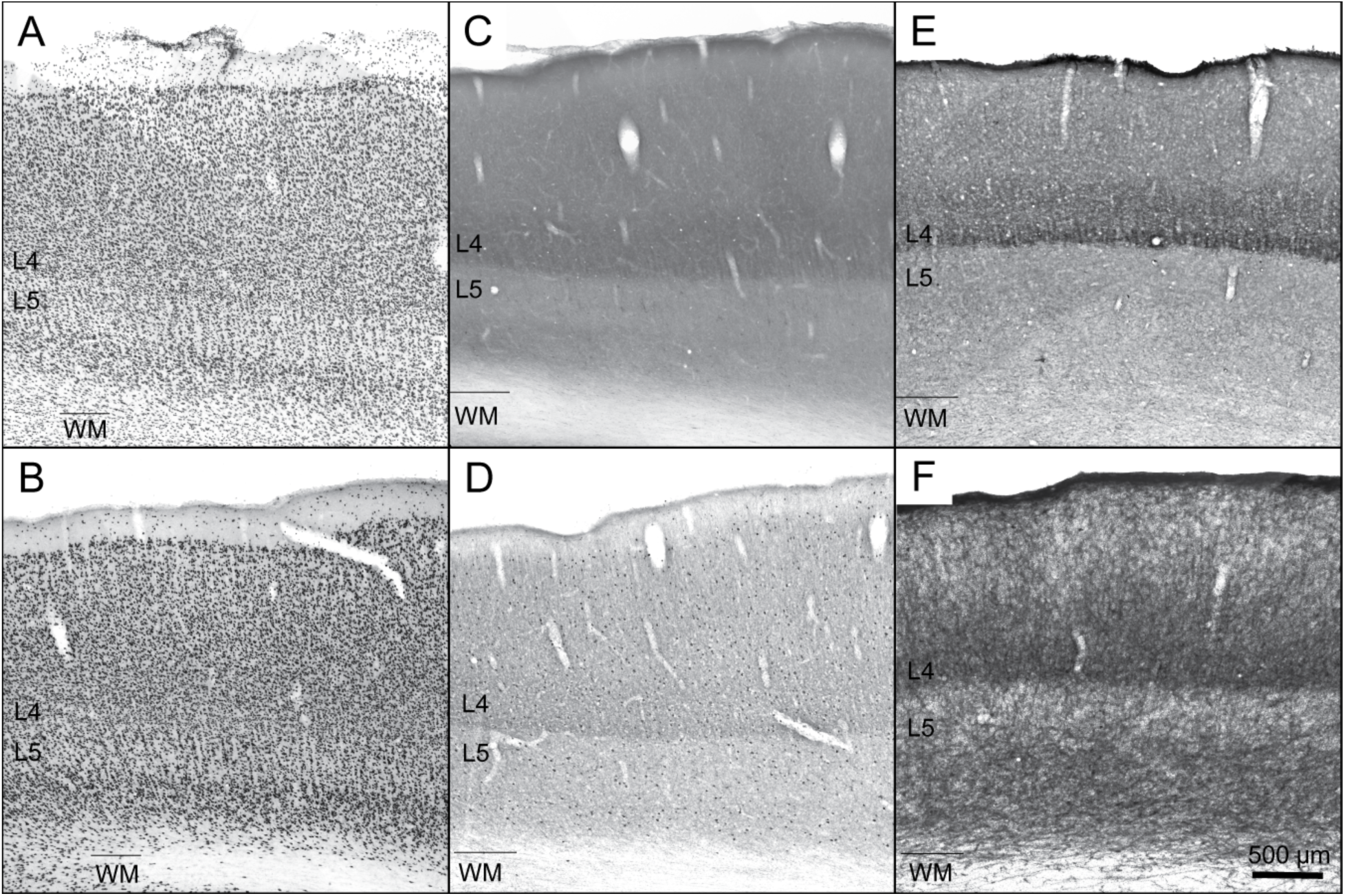
Characterization of layer 4 within V1 for *Aotus trivirgatus*. Low-magnification composite images (×10 objective) of adjacent coronal sections through V1 in the owl monkey (*Aotus trivirgatus*), illustrating the laminar position of the granular cell layer 4 (L4). (A) Nissl (thionin); (B) NeuN; (C) cytochrome oxidase; (D) parvalbumin; (E) VGLUT2; (F) acetylcholinesterase. L4, layer4, L5, layer 5, WM, white matter. Scale bar = 500 µm.

**Figure S4.**
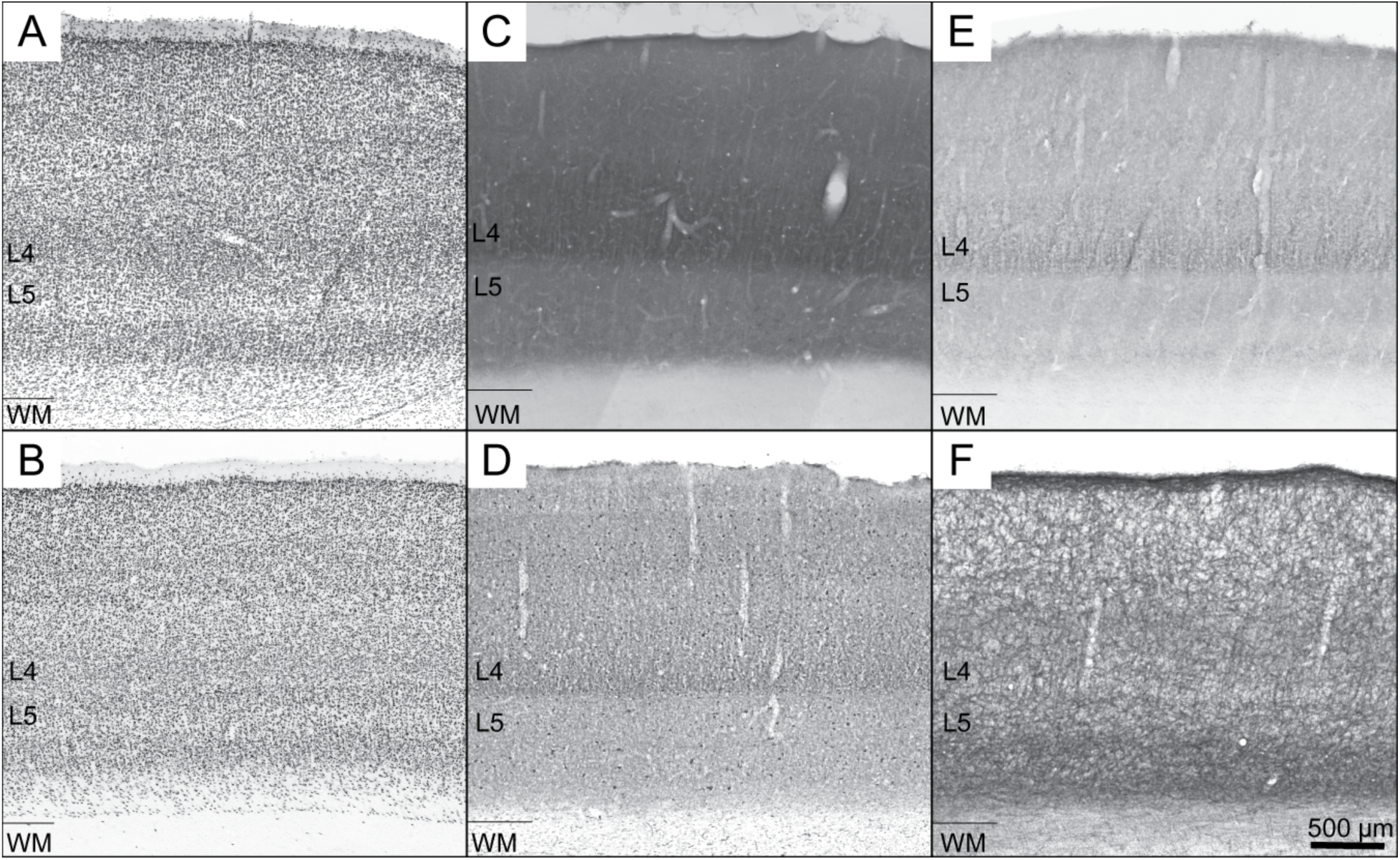
Characterization of layer 4 within V1 for *Otolemur garnettii*. Low-magnification composite images (×10 objective) of adjacent coronal sections through V1 in the prosimian galago (*Otolemur garnettii*), illustrating the laminar position of the granular cell layer 4 (L4). (A) Nissl (thionin); (B) NeuN; (C) cytochrome oxidase; (D) parvalbumin; (E) VGLUT2; (F) acetylcholinesterase. L4, layer4, L5, layer 5, WM, white matter. Scale bar = 500 µm.

**Figure S5.**
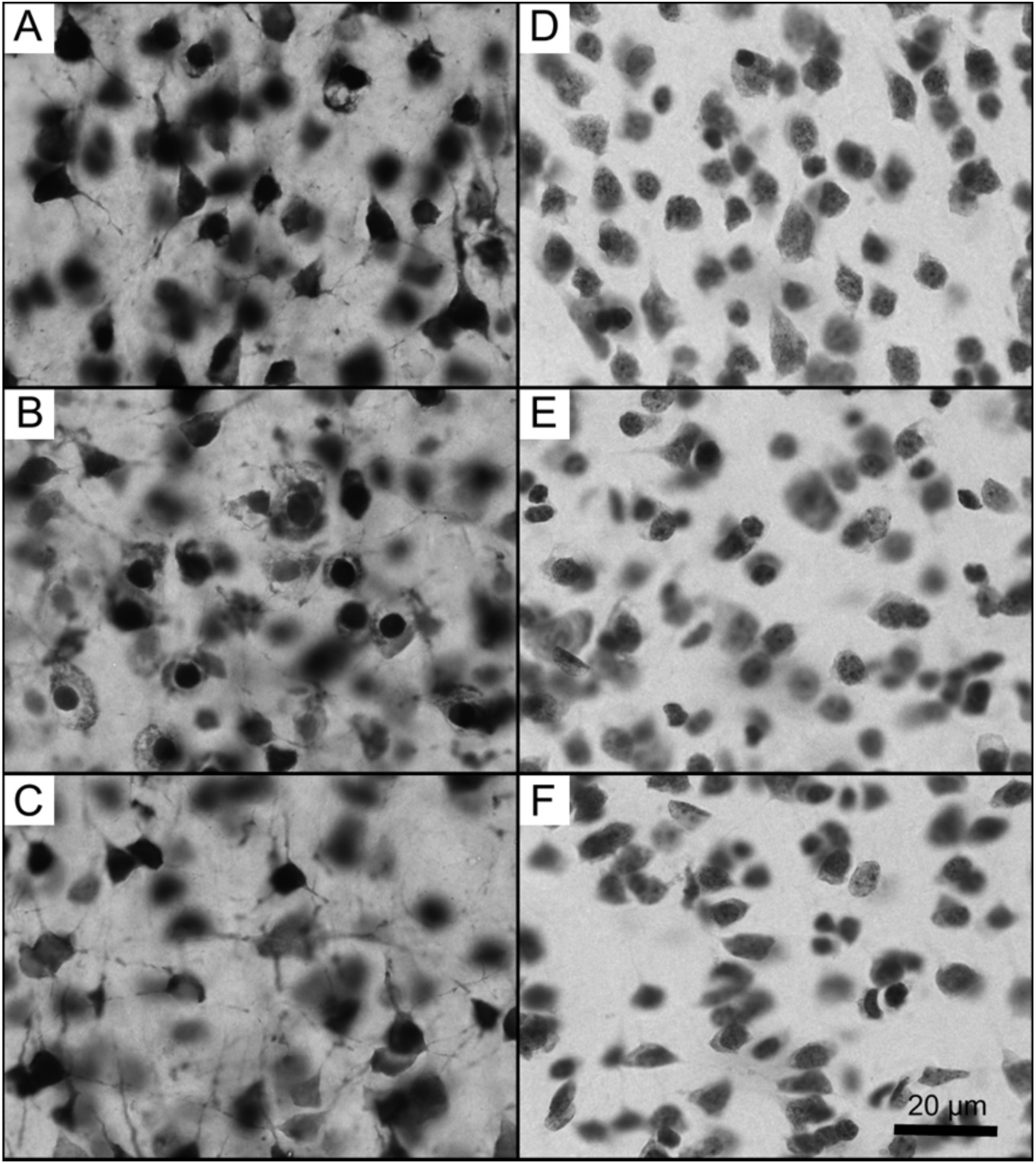
High magnification images of V1 cells in the macaque monkey (*Macaca mulatta*). High-magnification photomicrographs of V1 in the macaque monkey (*Macaca mulatta*), illustrating the cellular morphology and packing density used to distinguish neurons from glia in stereological analyses. The left panels (A–C) are sections immunolabeled for neuronal nuclei (NeuN), and the right panels stained for Nissl substance (D–F). Top row (A, D) are supragranular, middle row (B, E) are granular or layer 4, and bottom row is infragranular (C, F) or inclusive of layer 5. Scale bar = 20 µm.

**Figure S6.**
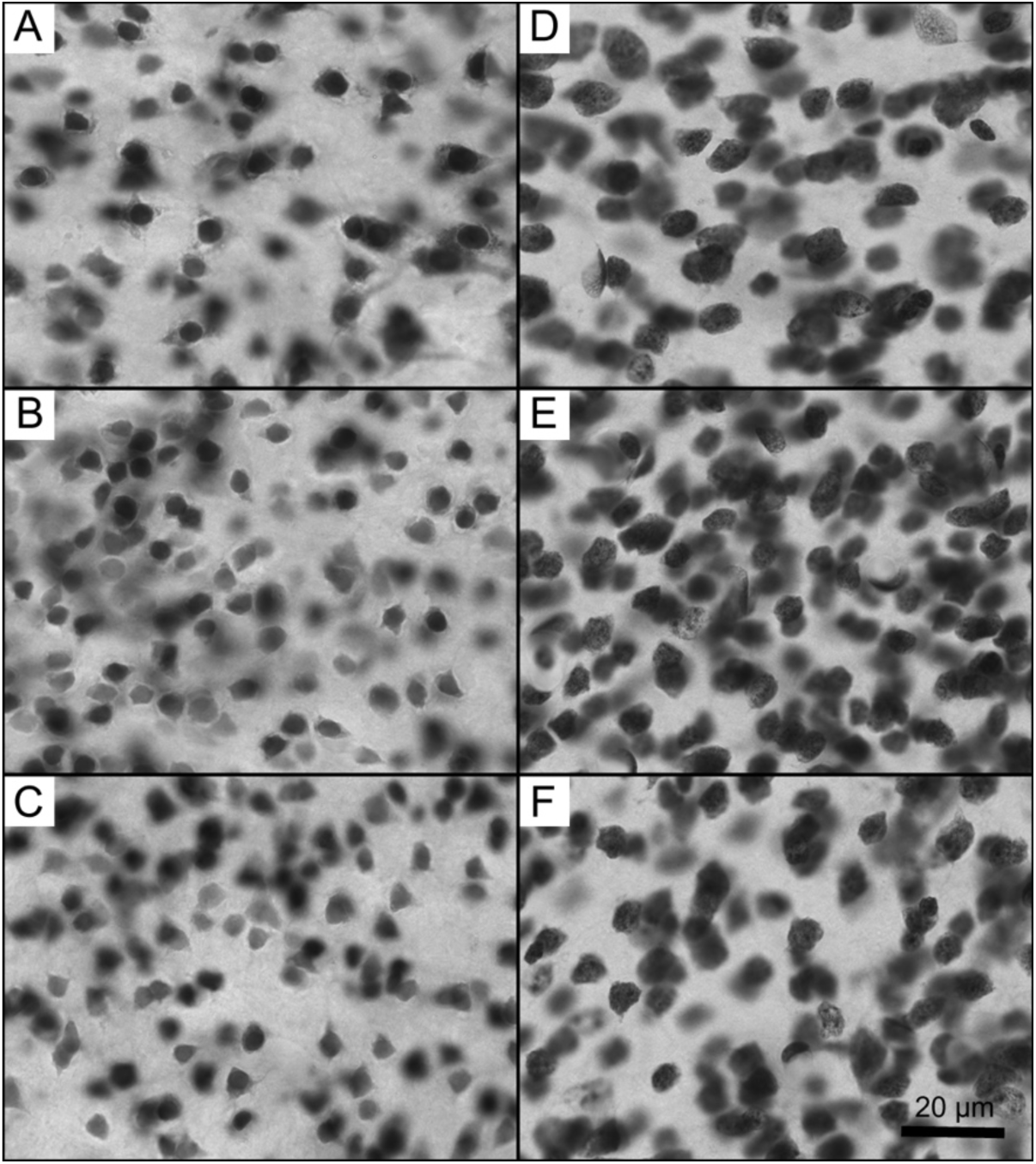
High magnification images of V1 cells in the squirrel monkey (*Saimiri sciureus*). High-magnification photomicrographs of V1 in the squirrel monkey (*Saimiri sciureus*). The left panels (A–C) are sections immunolabeled for neuronal nuclei (NeuN), and the right panels stained for Nissl substance (D–F). Top row (A, D) are supragranular, middle row (B, E) are granular or layer 4, and bottom row is infragranular (C, F) or inclusive of layer 5. Scale bar = 20 µm.

**Figure S7.**
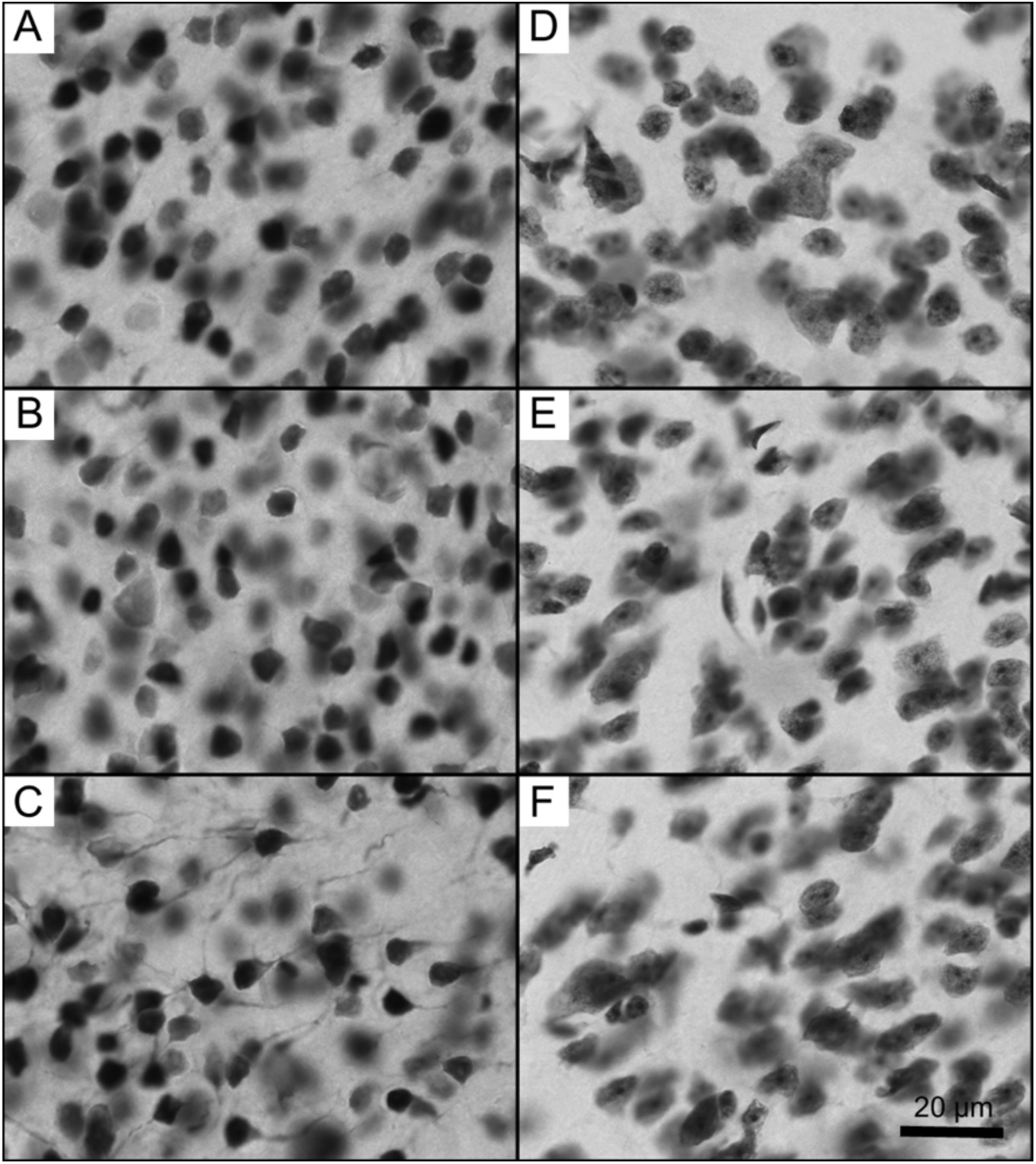
High magnification images of V1 cells in the owl monkey (*Aotus trivirgatus*). High-magnification photomicrographs of V1 in the owl monkey (*Aotus trivirgatus*). The left panels (A–C) are sections immunolabeled for neuronal nuclei (NeuN), and the right panels stained for Nissl substance (D–F). Top row (A, D) are supragranular, middle row (B, E) are granular or layer 4, and bottom row is infragranular (C, F) or inclusive of layer 5. Scale bar = 20 µm.

**Figure S8.**
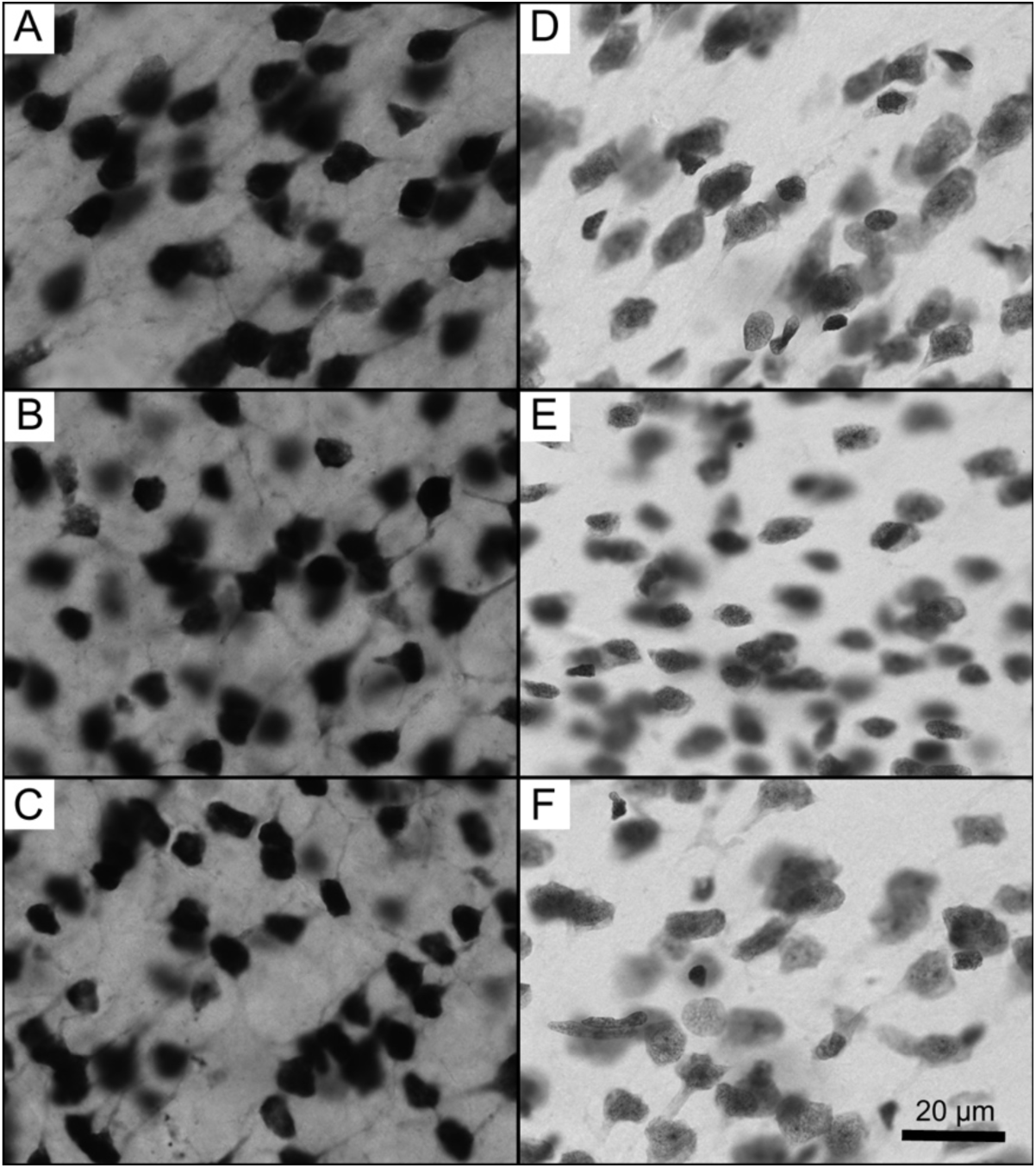
High magnification images of V1 cells in the galago (*Otolemur garnettii*). High-magnification photomicrographs of V1 in the prosimian galago (*Otolemur garnettii*). The left panels (A–C) are sections immunolabeled for neuronal nuclei (NeuN), and the right panels stained for Nissl substance (D–F). Top row (A, D) are supragranular, middle row (B, E) are granular or layer 4, and bottom row is infragranular (C, F) or inclusive of layer 5. Scale bar = 20 µm.

**Figure S9.**
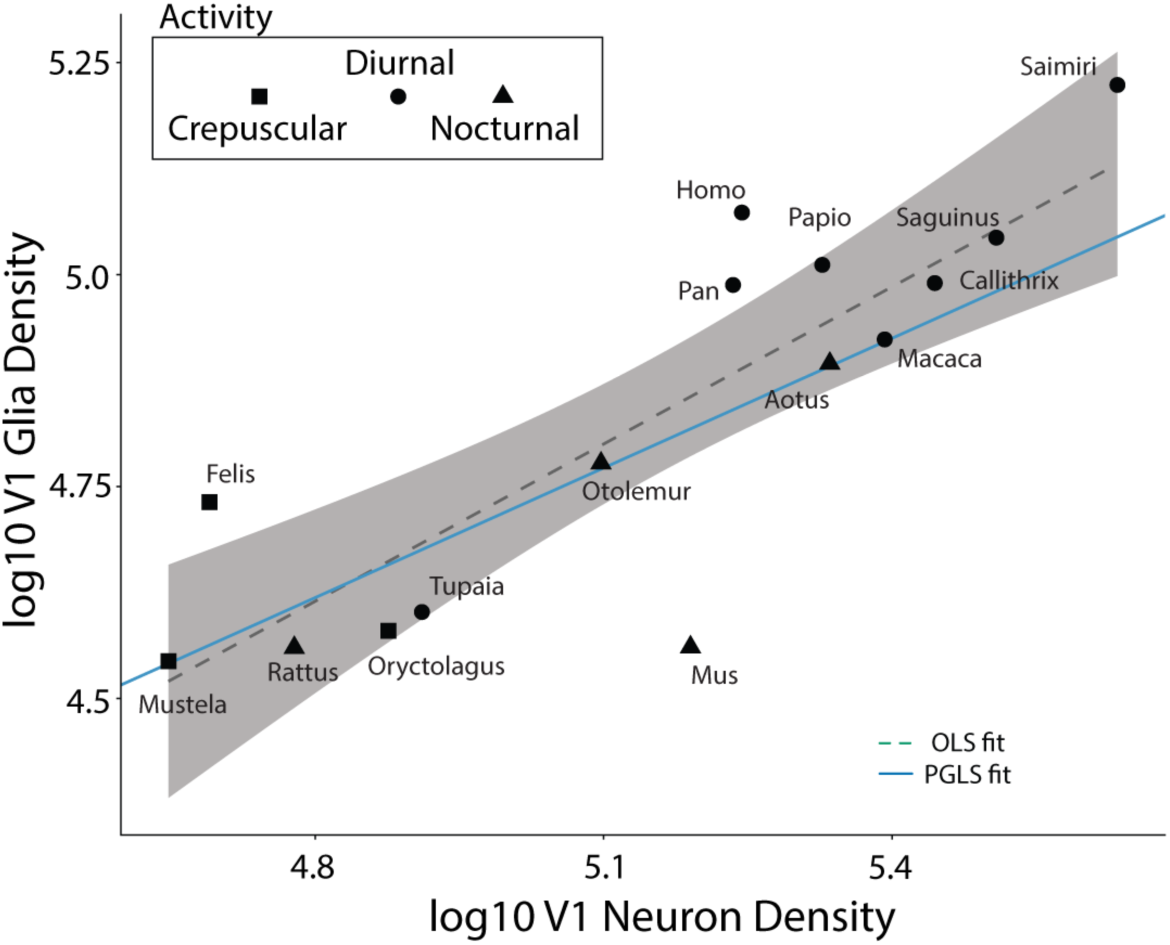
Scatterplot of log glia against neuron densities in V1. Log₁₀-transformed V1 glia density (neurons/mm³; y-axis) plotted against log₁₀-transformed V1 neuron density (neurons/mm³; x-axis) for all 15 species in the comparative sample spanning five mammalian orders (Primates, Rodentia, Carnivora, Lagomorpha, Scandentia). The dashed line and grey shading show the ordinary least squares (OLS) regression fit with 95% confidence interval; the solid blue line shows the phylogenetic generalized least squares (PGLS) fit. Symbol shape indicates activity pattern (circles, diurnal; triangles, nocturnal; squares, crepuscular) and symbol color indicates clade membership (legend below). Each data point is labeled by genus name.

**Figure S10.**
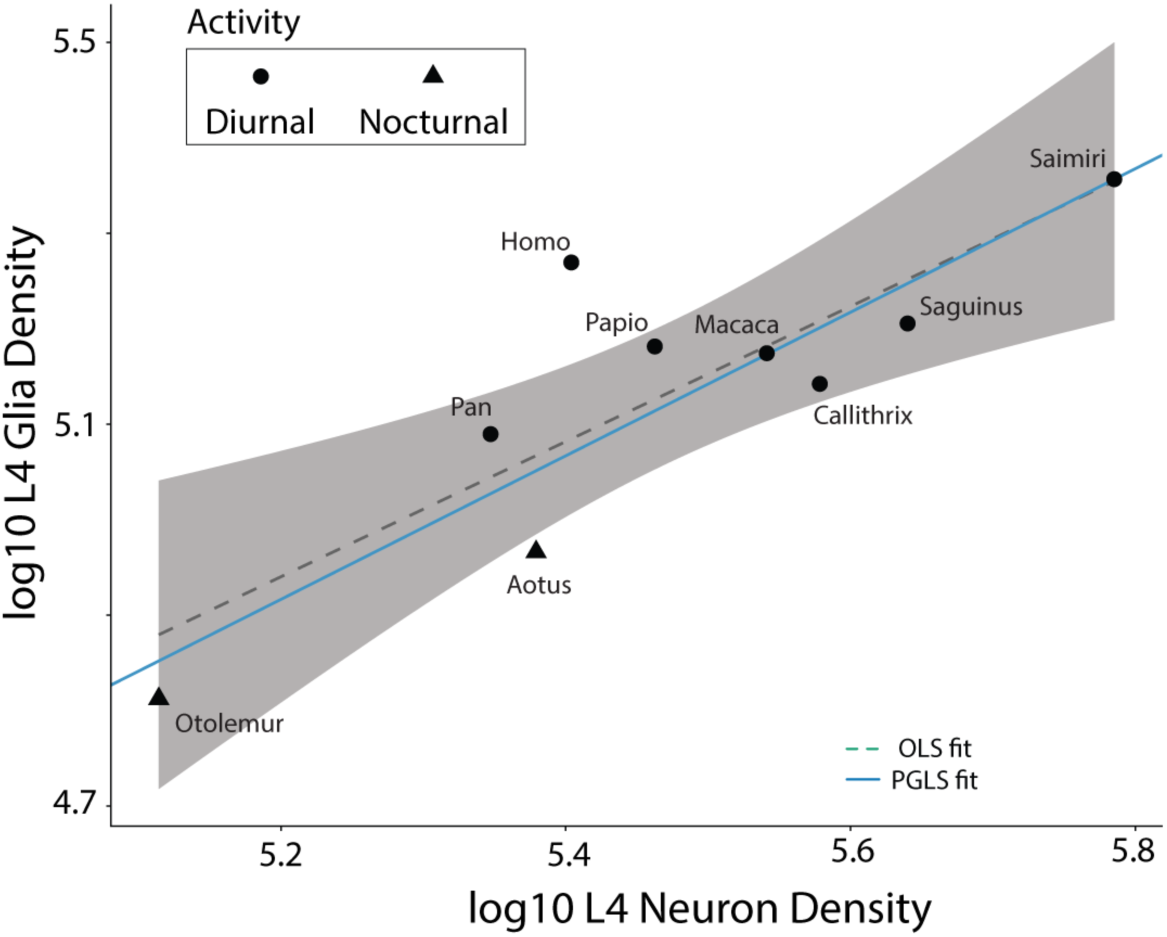
Scatterplot of glia against neuron densities in layer 4. Log₁₀-transformed layer 4 glia density (glia/mm³; y-axis) plotted against log₁₀-transformed layer 4 neuron density (neurons/mm³; x-axis) for the nine primate genera for which original stereological layer 4 estimates were obtained in this study. The dashed line and grey shading show the OLS regression fit with 95% confidence interval; the solid blue line shows the PGLS fit. Symbol shape indicates activity pattern (circles, diurnal; triangles, nocturnal) and symbol color indicates clade membership (legend below). Each data point is labeled by genus name. The near-isometric slope (PGLS slope = 0.95, Pagel’s λ ≈ 1.0) indicates that glia and neuron densities covary proportionally within layer 4, and that this covariation closely follows phylogenetic expectations under Brownian motion. *Homo* sits above both regression lines, consistent with the elevated layer 4 GNR (0.734) reported in the main text.

**Figure S11.**
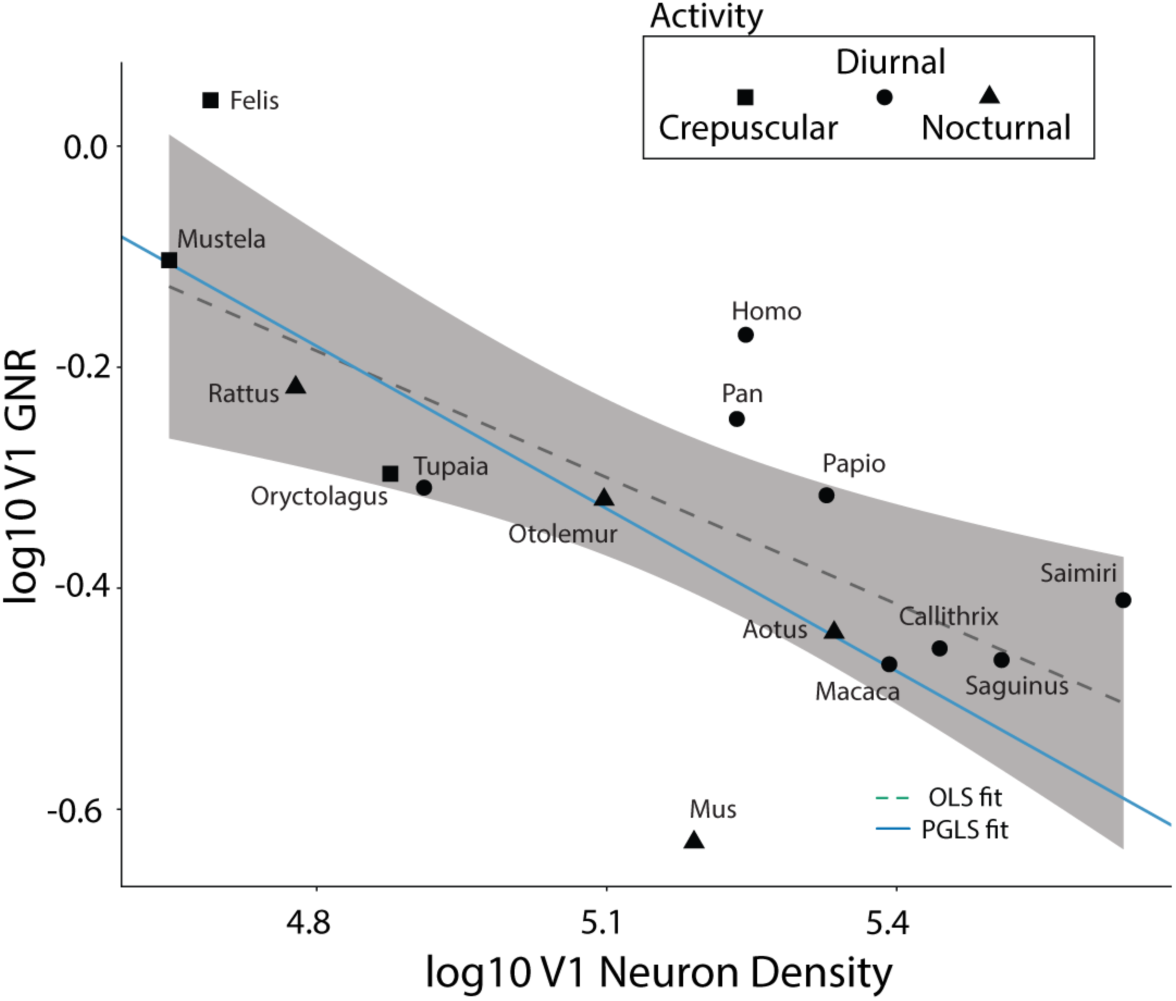
Scatterplot of GNR against neuron density. Log₁₀-transformed V1 glia-to-neuron ratio (GNR; y-axis) plotted against log₁₀-transformed V1 neuron density (neurons/mm³; x-axis) for all 15 species in the comparative sample spanning five mammalian orders. The dashed line and grey shading show the OLS regression fit with 95% confidence interval; the solid blue line shows the PGLS fit (slope = −0.490, SE = 0.143, p = 0.005, R² = 0.473, Pagel’s λ = 0.617). Symbol shape indicates activity pattern (circles, diurnal; triangles, nocturnal; squares, crepuscular) and symbol color indicates clade membership (legend below). Each data point is labeled by genus name.

**Figure S12.**
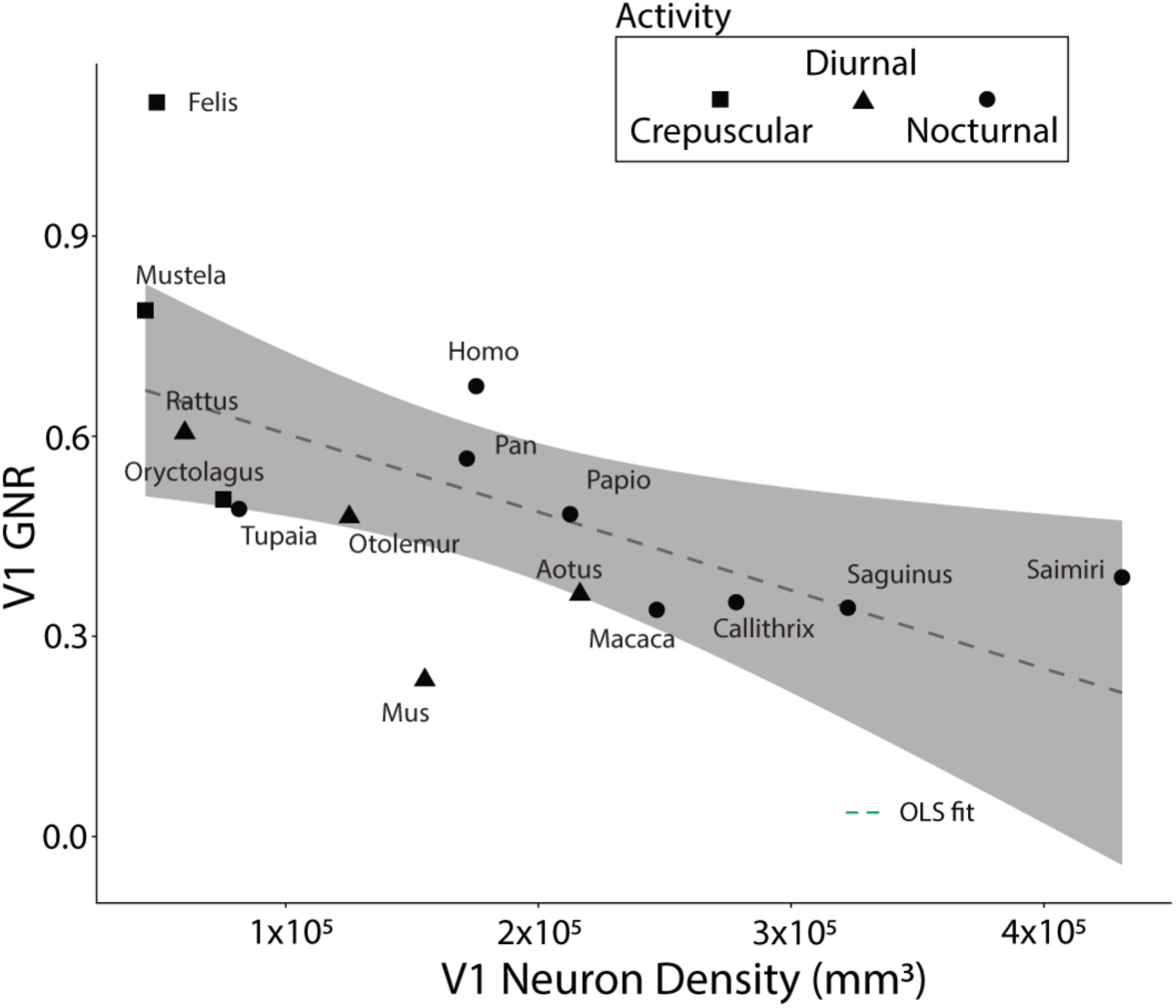
Scatterplot of GNR against neuron density. V1 glia-to-neuron ratio (GNR; y-axis) plotted against V1 neuron density (neurons/mm³; x-axis) on untransformed axes for all 15 species in the comparative sample spanning five mammalian orders. The dashed line and grey shading show the OLS regression fit with 95% confidence interval; no PGLS overlay is shown on raw-scale axes. Symbol shape indicates activity pattern (circles, diurnal; triangles, nocturnal; squares, crepuscular) and symbol color indicates clade membership. Each data point is labeled by genus name. This panel presents the same relationship as Supplemental Figure SF11 on arithmetic rather than logarithmic axes; statistical parameters are reported for the log-transformed model in Table 2 and Table S2. *Felis* occupies an outlier position at the upper left owing to its comparatively low neuron density and high GNR relative to the regression trend; *Homo* falls above the regression at moderate neuron density, consistent with the elevated whole-mantle GNR (0.675) reported in the main text.

**Dataset S1.** Comparative dataset on the mammal visual system.

**Dataset S2.** Visual circuit organization and amplification in mammals.

**Dataset S3.** Energy perturbation analysis.

**Dataset S4.** Full statistics on phylogenetic generalized least squares.

**Dataset S5.** Posterior trees robustness analysis.

**Dataset S6.** Jackknife leave-one-out analysis.

**Dataset S7.** Ordinary versus phylogenetic generalized least squares comparison.

**Dataset S8.** Longevity model series analysis.

